# The gut insulin receptor acts as gatekeeper of epithelial barrier integrity

**DOI:** 10.1101/2025.09.29.677561

**Authors:** Florian Sicherre, Dalale Gueddouri, Michèle Caüzac, Adèle Caudron-Hedeline, Fadila Benhamed, Wafa Charifi, Benoît Terris, Benoit Chassaing, Lucie Beaudoin, Matthieu Rouland, Angela Bosh, Claudia Cavelti-Weder, Gunter Leuckx, Yves Heremans, Agnès Lehuen, Catherine Postic, Gaëlle Boudry, Anne-Francoise Burnol, Sandra Guilmeau

## Abstract

An impaired gut barrier has emerged as a potential driver of the low-grade inflammation that accompanies obesity and its complications. Among these, hyperglycemia *per se* has been proposed to sustain such increased intestinal permeability and subsequent translocation of bacterial endotoxins in the systemic circulation. Because reduced insulin signaling in the gut epithelium has also been reported upon obese conditions, we hypothesized that, beyond hyperglycemia, defective intestinal insulin signaling could directly compromise epithelial integrity. To mimic this diabesity feature, we induced deletion of the insulin receptor (IR) in the adult gut epithelium of IR^ΔGUT^ mice. Remarkably, gut IR loss persistently maintained normal body weight and glucose homeostasis, thereby allowing the specific role of insulin action to be investigated. While IR^ΔGUT^ mice exhibited increased intestinal paracellular permeability, mechanistic characterization of this gut leakiness revealed that IR^ΔGUT^ mice displayed a rapid and drastic decline in Paneth cells anti-microbial defenses. This paralleled the onset of a cecal dysbiosis, as characterized by increased abundance of *Pseudomonadota*, and enhanced microbiota encroachment. Of note, IR^ΔGUT^ mice exhibited intestinal stem cell (ISC) defects, as evidenced by reduced expression of ISC markers and ISC-mediated growth of intestinal organoids. Although expression of niche factors such as *Wnt3a* was diminished in Paneth cells isolated from IR^ΔGUT^ mice, pharmacological activation of the canonical Wnt pathway failed to rescue the growth defects of IR-deleted gut organoids. The direct contribution of IR-downstream signaling to ISCs homeostasis was confirmed by the transcriptional reprogramming of FACS-sorted ISCs from IR^ΔISC^ mice. Finally, while gut IR loss did not worsen endotoxemia or impaired glycemic control upon HFD-feeding, IR^ΔGUT^ displayed a higher susceptibility to chemically induced colitis and enteric infections (*S. typhimurium*, *C. rodentium*), underscoring intestinal insulin signaling as a key determinant of barrier integrity and epithelial homeostasis.

## INTRODUCTION

Besides being the largest host-environment interface, the intestine is exposed to the highest microbial and dietary antigen load. To regulate interactions between luminal factors and the local immune system, the intestinal mucosa forms a selective barrier, that ingeniously allows the efficient transcellular transport of nutrients while strictly preventing the paracellular flow of immune-stimulatory bacterial products. The physical component of the gut barrier consists of a monolayer of rapidly renewing epithelial cells, sealed by intercellular junctions and overlaid with a mucus layer secreted by goblet cells. Antimicrobial peptides derived from Paneth cells, together with immune cells scattered beneath the epithelial layer, provide an additional chemical defense against enteric pathogens while maintaining tolerance to commensal bacteria.

Barrier disruption can lead to a « leaky gut », and the ensuing immune activation is pivotal in the pathogenesis of multiple diseases. Beyond gastrointestinal disorders, gut leakiness has emerged as a potential driver of the low-grade inflammation that accompanies metabolic alterations upon obese and diabetic conditions. Indeed, a common feature of these pathologies is their association with chronic inflammatory processes in various tissues, as well as an increased general risk of infection ^1–3^. A persistent positive energy balance indeed leads to a subclinical inflammatory state, which is considered to pave the way for insulin resistance and subsequent type 2 diabetes (T2D). Alongside other drivers, impaired intestinal mucosal barrier, and subsequent translocation of microbial by-products into the systemic circulation, are thought to contribute to this state of metaflammation ^4^. While alterations in gut barrier components have been proposed as an early event in the development of obesity ^5,6^, major gaps remain to be elucidated in mapping the mechanisms that elicit or sustain defective epithelial integrity upon the diabesity cascade. Multiple evidence have indicated that microbial imbalance, dietary components (e.g., high-fat diet, HFD) and hyperglycemia as prime candidates ^4,7,8^. Yet, because most studies have focused on obese and/or diabetic models, disentangling the specific contributions of adipose tissue remodeling, insulin resistance, and hyperglycemia to gut leakiness has proven challenging. In this context, our group recently evidenced that, independently of obesity, S961-induced insulin resistance impairs epithelial gut integrity in lean mice ^9^. Moreover, restoration of barrier function in mice lacking intestinal GLUT2 or following inhibition of glucose metabolism has demonstrated that hyperglycemia *per se* disrupts gut integrity via retrograde glucose flux ^7^.

Because hyperglycemia and loss of insulin action are often two sides of the same coin, we hypothesized that, beyond hyperglycemia, defective intestinal insulin signaling *per se* could directly impair epithelial integrity. While the gut has long been recognized as an insulin-sensitive organ, the specific detection of insulin receptor (IR) in intestinal epithelial cells opened new avenues regarding the direct roles for insulin in the control of gut mucosa homeostasis ^10^. Of note, impaired insulin signaling was reported in the intestinal epithelium of HFD-induced obese mice and in jejunal explants from obese subjects ^11,12^, pointing to defective insulin action in such a pathological state. Previous studies using constitutive mouse models with gut IR deficiency ^11–13^ reported modest phenotypes under both basal and HFD feeding conditions, such as reduced glucose absorption and decreased GIP incretin content in the gut epithelium ^13–15^. Because none of them clearly documented either the cellular or the functional consequences of impaired intestinal insulin signaling on intestinal barrier function, we developed a mouse model with an inducible deletion of the insulin receptor in the adult gut epithelium (IR^ΔGUT^), thereby mimicking a key feature of diabesity. Altogether, this study shows that lean, normoglycemic IR^ΔGUT^ mice exhibit intestinal hyper-permeability, indicating impaired epithelial insulin signaling as a novel candidate mechanism underlying gut leakiness associated with metabolic syndrome. Accordingly, we provide evidence that direct epithelial insulin action controls two essential components of the overall gut barrier function, *i.e.* antimicrobial defense by Paneth cells and intestinal stem-cell dynamics. Finally, loss of intestinal IR sensitizes the host to both inflammatory and infectious injuries, underscoring the protective role of insulin signaling in maintaining intestinal homeostasis.

## MATERIAL & METHODS

### Animals and treatments

All procedures were carried out according to the French guidelines for the care and use of experimental animals and were approved by the “Direction départementale des services vétérinaires de Paris” with the following APAFIS numbers: # 14856-2018041615405803; #15482-2018041615405803; #15573-2018072014264041; #19178-201902131234408. Mice carrying the LoxP sites flanking the fourth exon of the *insr* gene, that encodes the insulin receptor (IR^lox/lox^ mice, stock number: 006955, Jackson Laboratory, USA), were crossed with C57BL/6J, which express an inducible Cre recombinase driven by the 9-kb mouse villin1 promoter (Vil-Cre^ERT2^ mice, kind gift from Dr Sylvie Robine) ^15^. The resulting IR^lox/+; Vil-CreERT2^ mice were interbred with IR^lox/lox^ mice to generate IR^lox/lox; Vil-CreERT2^ mice, referred as IR^ΔGUT^ (for inducible gut insulin receptor knock-out). IR^lox/lox^ mice were also crossed with Lgr5-EGFP-IRES-CreERT2 transgenic mice (kindly provided by Dr. Béatrice Romagnolo) to generate IR^lox/lox; Lgr5-EGFP-IRES-CreERT2^ referred as IR^ΔISC^ (for inducible intestinal stem cell insulin receptor knockout). All mice were housed in colony cages with a 12-h light/dark cycle in a temperature-controlled environment and had free access to water and regular diet (65% carbohydrate, 11% fat and 24% protein). Ten-week-old male IR^fl/fl^ and IR^ΔGUT^ mice were housed in separate cages before induction of *insr* deletion by tamoxifen (TMX) and were sacrificed 3 days (short term) or 10 weeks (long term) following the beginning of the treatment. IR^fl/fl^ and IR^ΔGUT^ mice were orally administered (gavage) TMX citrate salt (1 mg/ mouse/ day, Sigma T9262) diluted in 5% carboxymethylcellulose sodium salt (Sigma) during 5 consecutive days, except for the short-term study, where mice were treated for only 3 days. For IR^ΔISC^ mice, *insr* deletion was induced by TMX intraperitoneal injection (1 mg/ mouse/ day; MP Biomedicals 156738) dissolved in food-grade sunflower oil during 4 consecutive days. Comparative evaluation of the metabolic profiles of IR^fl/fl^ and IR^ΔGUT^ was performed by feeding these two groups with either a chow diet or a coconut-based high fat diet HFD (60%; #D12331, Research Diets or Sniff) for 12 weeks.

### Oral glucose and insulin tolerance tests

Oral glucose tolerance tests (OGTT) were performed after overnight fasting. Mice received a glucose solution administered either by oral gavage (2 g/kg body weight, Sigma) or by intraperitoneal injection (IPGTT; 2 g/kg body weight, Braun, Germany). The blood glucose was measure at baseline and indicated timepoints post-injection or -gavage. For insulin tolerance tests (ITT), mice were fasted for 6 hours and injected ip with insulin (0.75 U/kg; Actrapid Penfill, Novo Nordisk). Blood glucose concentrations were measured at the indicated time points using an Accu-Chek glucometer (Roche Diabetes Care). Insulin levels were measured at baseline, 15 and 30 min using a mouse/rat insulin kit (MesoScale Diagnostics, USA).

### Intestinal permeability

Mice were submitted to an oral gavage of FITC-dextran 4 kDa (FD4, 0.6 mg/g body weight) (Sigma 46944) and 0.5% carboxymethycellulose solution including 6% Carmine red (Sigma C5678) in order to evaluate gut transepithelial permeability normalized to intestinal transit. One hour after FD4 gavage, peripheral blood was collected and serum FITC-dextran concentration was determined using a fluorescence spectrophotometer (Ex/Em 490/520 nm) and further normalized to the transit time. The total intestinal transit time was defined as the interval between ingestion of carmine red and the first appearance of the dye in feces.

### Antibiotics treatment

Mice were treated for 5 days with antibiotics that were added sequentially in drinking water, as followed. Day 1: colistin (1 mg/ml) (Cayman chemical), day 2: colistin and vanomycin (45 μg/l) (Sigma), day 3: colistin, vanomycin and streptomycin (1 g/l) (Sigma), day 4: colistin, vanomycin, streptomycin and ampicillin (1 mg/ml) (Sigma). Caecal content was collected for further bacterial DNA analysis (Quick-DNA fecal miniprep Kit (Zymo research).

### Enteropathogenic bacterial infections

For *Citrobacter rodentium* (strain ICC180, expressing luciferase) infection, the inoculum was prepared as previously described ^18^, and an oral gavage of 10^9^ cfu bacteria per mouse (200 µl suspension in PBS) was performed. Bacterial colonization was assessed by kinetic analysis of bioluminescence using *in vivo* imaging (IVIS System) and by daily fecal sample collection. For *Salmonella Typhimurium* infection (strain SL1344), mice were pre-treated with streptomycin (20 mg by oral administration, Sigma) after a 4 h fast. The following day, mice were fasted for 4 hours before oral gavage with 2×10^8^ cfu (day 0). Body weight was further measured daily and fecal samples were collected over 4 consecutive days. Serial dilutions from various organ homogenates were plated onto Luria-Bertani (LB) agar plates containing 50 µg/ml streptomycin, and colony counts were performed after overnight incubation at 37°C.

### DSS-induced colitis

Colitis was induced by adding 2% Dextran Sulfate Sodium (DSS, mpbio 160110) in drinking water in 3- to 4-month-old male mice the week following the TMX gavage ^19^. After 5 days, DSS was removed and tap water was given back to the mice. Body weight was measured daily and mice were sacrificed at 8 to 10 days after beginning of the DSS challenge. At sacrifice, colon was collected from each mouse for further analyses.

### Gut organoids cultures

The USI was opened longitudinally, and incubated on ice for 30 min in 15 mM EDTA-PBS. Enriched crypt fractions after 6 to 8 cycles of vigorous vortexing (30 s) and EDTA incubation, and further filtration of the suspension through a 70 µm cell strainer and centrifugation at 250 g for 5 min. The resulting crypt pellet was resuspended and plated in Matrigel (Corning #356231) at approximately 350 crypts per well in 24-well plates. Organoid cultures were then maintained in ENR medium ^16^ that contained DMEM/F12 medium (ThermoFisher Scientific #12634010), 10 mM HEPES (ThermoFisher Scientific) supplemented with antibiotics (ThermoFisher Scientific) and 500 ng/ml R-spondin-1 (R&D systems #3474-RS), 100ng/ml RNoggin (Peprotech #250-38) 50ng/ml EGF (Peprotech #315-09), N2-supplement 1X and B-27 Supplement 1 X (Life-technologies #17502048 and #17504044). As already published ^17^, 1 mM of valproic acid and/or 3 µM of CHIR99021 (Tocris #4423) were added the ENR medium for ENR-CV and ENR-C conditions respectively (Sigma #P4543). Organoid growth was then monitored by Incucyte scanning every 4 h at a 4× magnification. Image analysis parameters included: radius 125 µm; sensitivity 20; edge split enabled; edge sensitivity 50; hole fill 5×10⁵ µm²; minimum area 1200 µm²; maximum eccentricity 0.95.

For insulin challenge experiments, intestinal crypts were seeded in insulin-free ENR medium formulated with DMEM/F12 GlutaMAX (ThermoFisher Scientific #31331028) and without N2-supplement, that contains insulin. Twelve hours post-seeding, organoids were treated with 100 nM insulin for 5min, after which the medium was removed and organoids were immediately snap-frozen for further Western blot analyses.

### In vitro irradiation assay

Gut organoids were cultured for 3 days, then imaged at 4× magnification (white field, Z-step 120 µm) in a 37 °C chamber. Plates were then individually exposed to 4 Gy irradiation. Imaging was repeated at days 3 and 6 post-irradiation using the same conditions.

### In vitro MTT assay

After addition of 500 µg/mL MTT solution to the culture medium for 2 h at 37 °C, the medium was removed and Matrigel was solubilized in 100 µL of 2% SDS, followed by the addition of 515 µL DMSO. The solution was homogenized 1 h after the addition of DMSO, and 200 µL aliquots were transferred to a 96-well clear bottom plate for absorbance reading at 562 nm on a CLARIOstar reader (BMG Labtech). Blanks contained SDS + DMSO without organoids. Assays were conducted at days 1, 3, and 5 post-seeding.

### Gut epithelial cells and crypts isolation

Intestinal epithelial cells (IECs) were isolated from both the upper and the lower small intestine (USI: duodenum and jejunum, LSI: ileum). Tissues were initially incubated for 5 minutes at 37 °C in a washing solution (1.5 mM KCl, 96 mM NaCl, 27 mM sodium citrate, 8 mM KH₂PO₄, 5.6 mM Na₂HPO₄, 1 mM DTT). Samples were then transferred into a chelating buffer (15 mM EDTA, 1 mM DTT, and one protease inhibitor tablet per 500 mL PBS; Roche) and incubated for 30 min at 37 °C under gentle agitation (100 rpm). Isolated cells were centrifuged, flash-frozen in liquid nitrogen, and stored at −80 °C until further RNA and protein analyses. In order to prepare crypt-villus fractions from the small intestine, villi were after the washing solution isolated in 1.5 mM EDTA in PBS for 25 min, and the remaining material was treated as described above for isolated IEC (15 mM EDTA in PBS for 30 min).

### Flow cytometry

Regarding DSS challenge experiments, colonic cells from the *lamina propria* were isolated using Lamina Propria Dissociation Kit (#130-097-410, Miltenyi) as previously described ^18^. Each subset of *lamina propria* cells was analyzed via flow cytometry after 20 min of staining at 4°C in PBS containing 2% fetal bovine serum and 0.1% sodium azide with the following antibodies: anti-CD45 (APC-Cy7, Biolegend 30F11), anti-CD64 (BV605, X54-57.1 Biolegend), anti-Ly-6C (BV510, HK1.4 Biolegend), anti-Ly-6G (PE, 1A8 BD Biosciences), anti-CD11b (BV785, M1/70 Biolegend), anti-CD11c (APC, N418 Invitrogen), anti-CD103 (BV421, M290 BD Biosciences), anti-CX3CR1 (PE-CF594, SA011F11 Biolegend), anti-F4/80 (PE-Cy7, BM8 Biolegend), and anti-CMH ClassII (AF700, M5/114-5-2 Invitrogen). Data acquisition was processed using Biosciences LSR Fortessa (BD Biosciences). Gating and calculation of cell population frequencies were computed with FlowJo (Tree Star) analysis software. Macrophages were identified as CD45^+^ <u>CD19</u>^-^ TCRγδ^-^TCRβ^-^ CD11b^+^ cells, DCs were identified as CD45^+^ CD19^-^ TCRγδ^-^ TCRβ^-^ CD11b CD206^-^ CD11c^+^ cells, T cells were identified as CD45^+^ CD19^-^ TCRβ^+^ cells, and B cells were identified as CD45^+^ TCRβ^-^ CD19^+^ cells. Gating strategies to determine macrophages, DC, T, and B cells were as previously described ^9^.

For Paneth cell sorting, small intestinal crypts were dissociated into single cells using pipetting in a solution containing Y-27632 (Enzo LifeScience ALX-270-333; 5 mM), dispase (Corning, 6U/mL), and DNase I (ThermoFisher Scientific; 8U/µl), incubated at 37 °C until complete dissociation. Cell suspensions were filtered through 40 µm strainers, pelleted (500 g, 5 min, 4 °C), then resuspended in dissociation buffer without enzymes. After counting with a TC20 Automated cell counter (Biored), cells were preincubated with FC-block (Biolegend 423105; 25 µg/10⁶ cells in 100 µL PBS) on ice for 10 min, followed by staining with antibody cocktails (quantity are given for 10⁶ cells in 100 µL PBS): anti-CD45-APC-Cye7 (Biolegend 103115; 62,5 ng), anti-CD31-APC-cye7 (Biolegend 102533; 62,5 ng), anti-Ep-CAM-BV785 (Biolegend 118245; 2,5ng), anti-CD24-PE-dazzle594 (Biolegend 101837; 62,5 ng), anti-c-Kit PerCP-eFluor 710 (Invitrogen 46-1171-82; 31,25 ng) and the Zombie live/dead prob (Biolegend 423105; 0,1 µL). Staining was performed for 30 min on ice in the dark, then cells were washed and resuspended in flow buffer (PBS + 1 % BSA, 5 mM EDTA, 5 mM Y-27632). Paneth cells (PC^+^) were further identified as CD45^-^ CD31^-^ Zombie^-^ EPCAM^+^ CD24^+^ckit^+^ cells.

For intestinal stem cell sorting, cells were dissociated as for Paneth cells, stained with CD24-APC (6 µL per 10⁶ cells in 1 mL PBS + 1 % BSA, 10 µM Y-27632) for 20 min on ice, washed, and resuspended in flow buffer. Propidium iodide (PI, 1 µL) was added 5 min before FACS. Intestinal stem cells were further identified as PI^-^ GFP^+^ CD24^+^ cells.

### Biochemical analyses

Fecal lysozyme activity (EnzChek lysozyme assay kit, Molecular Probes), lipocaline-2 (Mouse Lipocalin-2/NGA Duo set ELISA kit, R&D Systems) MPO contents (Mouse MPO ELISA kit (HycultBiotech) as well as plasma LBP (mouse LBP ELISA kit, Abcam) and insulin (mouse ultrasensible insulin ELISA, ALPCO) concentrations were measured using the indicated commercial assays according to the manufacturers’ instructions.

### Western blot analyses

IECs were solubilized as previously described ^17^. The protein extracts were subjected to SDS-PAGE electrophoresis and immunoblotted with the following antibodies, diluted as followed: anti-pAkt (Ser473) (CST #4060, Cell Signaling Technology; 1:1000), anti-Akt (CST #9272; 1:1000), anti-β-actin (CST #4970; 1:1000), anti-PCNA (CST #2586; 1:2000), anti-IRβ (SC #711, Santa Cruz Biotechnology; 1:1000), anti-GAPDH (SC #25778; 1:1000). The immunoreactive bands were revealed using the Clarity Western ECL Substrate (BIO-RAD). Chemiluminescence analyses were performed with the ChemiDOc MP Imaging System (BIO-RAD).

### In situ histological analyses

Samples from proximal USI, LSI and/or colon were fixed in a 4 % PFA, embedded in paraffin and sectioned (5 µm). Following deparaffinization and rehydration, sections were stained with hematoxylin–eosin (HE) or Alcian Blue using standard protocols.

To score either DSS- or enteropathogens-induced inflammation, histopathological analyses of small intestinal and/or colonic sections were conducted blinded, based on 6 criteria: inflammation severity, ulceration depth, ulcerative area, edema, vascular dilation, and ischemia. A cumulative “total histological score” was calculated for each sample.

For immunohistochemistry, tissue sections were permeabilized with TBS-Tween 0.1 %, blocked with 3 % BSA, then incubated with the following diluted primary antibodies: anti-Ki-67 (CST #9129; 1:300), anti-Lysozyme (Dako #EC.3.2.1.17; 1:500), or anti-Olfm4 (CST #39141; 1:400).

Co-immunofluorescence stainings were performed after blocking with 5 % BSA, using the following antibodies: primary anti-Lysozyme or anti-Olfm4 antibodies together with an anti-E-cadherin antibody (BD Biosciences #610081; 1:100), followed by secondary antibodies anti-rabbit and anti-mouse (respectively Alexa-Fluor 647 and Alexa-Fluor 555, Invitrogen; 1 :1000). Nuclei were counterstained with Vectashield and DAPI dye. Images were captured at 20× and 40× magnification.

The RNAscope Multiplex Fluorescent v2 Assay (# 323100, Advanced Cell Diagnostics) was used according to manufacturer’s instructions. Probes used were *B. subtilis* negative control *dapb* (# 310043), *M. musculus* positive control *ppib* (# 313911) and *M. musculus insr* (# 401011-C2), *Lgr5* (# 312171) followed by immunostaining for E-cadherin (# 610081 at 1:100, BD Biosciences), and anti-lysozyme (Dako, EC.3.2.1.17, 1:500). Imaging was performed with an Axio Scan.Z1 (Zeiss) at 40x magnification. RNA in situ hybridization, immunostaining, imaging and analysis was performed by the VSTA core facility at VUB (https://vsta.research.vub.be).

### Quantitation of bacterial encroachment

To analyze bacteria localization at the surface of the intestinal mucosa, mucus immunostaining was paired with fluorescent *in situ* hybridization, as previously described ^19^, Proximal colonic tissues containing fecal material were fixed in methanol-Carnoy solution (60% methanol, 30% chloroform, 10% glacial acetic acid) for 3 h at room temperature. Tissues were then washed twice in methanol (30 min), ethanol (15 min), ethanol/xylene (1:1, 15 min) and xylene (15 min), before paraffin embedding in a vertical orientation. Five μm sections were dewaxed at 60°C for 10 min, followed by 10 min incubation with xylene and 99.5% ethanol. Hybridization step was performed at 50°C overnight with EUB338 probe (5’-GCTGCCTCCCGTAGGAGT-3’, with a 5’ labeling using Alexa 647) diluted to a final concentration of 10 μg/mL in hybridization buffer (20 mM Tris–HCl, pH 7.4, 0.9 M NaCl, 0.1% sodium dodecyl sulfate, 20% formamide). After washing in wash buffer (20 mM Tris–HCl, pH 7.4, 0.9 M NaCl, 10 min) and twice in phosphate-buffered saline (PBS 10 min), tissue sections were blocked for 30 minutes at 4°C with 5% fetal bovine serum in PBS. After overnight incubation at 4°C with an anti-Mucin-2 antibody (SCT #SC15334; 1:1500), sections were washed 3 × 10 min in PBS, and further applied for 2 hours with an anti-rabbit Alexa 488 secondary antibody (1:1500), phalloidin-tetramethylrhodamine B isothiocyanate (Sigma, 1 μg/mL) and Hoechst 33258 (Sigma,10 μg/mL). After washing 3 × 10 minutes in PBS, sections were mounted using the Prolong antifade mounting media (Life Technologies). Observations of bacterial localization and quantitation of bacterial-epithelial distance were performed via confocal microscopy. Instrument software was used to determine the distance between bacteria and epithelial cell monolayer. For each sample, 5 high-powered fields (HPF) were arbitrarily selected with the following inclusion criteria: (1) the presence of stained bacteria, (2) the presence of a clear and delimitated mucosal layer, and (3) the presence of an intact mucus layer. For each HPF, the distance between the 5 closest bacteria and the epithelium was determined. Thus, each bacterial-epithelial distance indicated by a point in the figures is, in fact, the average distance of 25 bacteria-epithelial distances.

### Transmission electron microscopy

Samples from LSI were analyzed under a JEOL 1011 transmission electron microscope with a GATAN Erlangshen CCD (charge-coupled device) camera. Acquisitions were processed using software « Digital Micrograph » and analyzed on ImageJ.

### Quantitative RT-qPCR analyses

Total RNAs were extracted from isolated IECs and hepatocytes using QIAzol® (Qiagen) according to the manufacturer’s protocol and for each sample, 1 μg of total RNA was reverse-transcribed using a standard reverse transcriptase (Life Technologies). For intestinal organoids, culture medium was removed directly before adding QIAzol®, in which Matrigel domes were scraped and dissolved. To enhance RNA recovery of low-yield samples (FACS-sorted cell, organoids), RNA grade glycogen (ThermoFisher Scientific) was added during overnight precipitation of RNA at −20 °C. In case yield remained low, 100 ng total RNA was reverse-transcribed using SuperScript IV (Life Technologies). Quantitative PCR was performed on a LightCycler® 480 (Roche) using SYBR Green I Master mix, with indicated primers (Supplementary Table 1). Gene expression levels were then normalized using three housekeeping genes: *Gapdh (glyceraldehyde-3-phosphate dehydrogenase)*, *Tbp (ATA-box binding protein)*, and *Rn18s (18S rRNA)*.

### Transcriptomic analyses

For transcriptomic analyses of LSI crypts, RNAs were extracted from both IR^ΔGUT^ and IR^lox/lox^ mice following treatment with tamoxifen for 3 consecutive days and further analyzed using Affymetrix clariomS Mouse arrays (Gene Expression Omnibus (GEO) dataset GSE81387). Samples were normalized using the RMA algorithm (Bioconductor affy package) and differential expression was measured with moderated t-test (limma R package). Gene set enrichment analyses (GSEA) of the transcriptomic data were performed using gene sets from BioCarta and KEGG databases.

For transcriptomic analyses on ISC from IR^ΔISC^ mice and their control group (Lgr5-EGFP-IRES-CreERT2), ISC were FACS-isolated after 4 days of TMX injection. RNA integrity (RIN > 8) was confirmed using an Agilent 2100 Bioanalyzer (pico chip). Starting from 4 ng total RNA, libraries were generated using the NEBNext low-input RNA kit: cDNA was synthesized via oligo-dT and template switching, adapters were ligated, and libraries were amplified, purified, quantified (Qubit HS DNA assay), and checked for size distribution. Sequencing was performed on an Illumina NextSeq 500 (paired-end, 75 bp, V2 chemistry). Raw reads were quality-checked (FastQC v0.11.5) and demultiplexed using AOZAN (ENS, Paris). Reads were aligned to Ensembl release 101 using STAR v2.7.6a, then quantified with RSEM v1.3.1. Differential expression analyses were carried out in R (v3.6.3) via DESeq2 (v1.26.0), using a median-of-ratios normalization and filtering out genes with fewer than 10 reads in fewer than three samples. The Wald test with Benjamini-Hochberg correction (FDR < 0.05) was applied to identify differentially expressed genes. Scripts and parameters are available on GitHub (https://github.com/GENOM-IC-Cochin/RNA-Seq_analysis). GSEA was subsequently performed on R using fgsea and clusterProfiler packages, leveraging MSigDB v7.0 gene sets.

### Microbiota 16 sequencing

Bacterial DNA was extracted from luminal content samples using the ZR fecal DNA Miniprep kit (Zymo Research, USA). The V3-V4 region of 16S rDNA was amplified using the following primers: CTTTCCCTACACGACGCTCTTCCGATCTACTCCTA-CGGGAGGCAGCAG (V3F) and GGAGTTCAGACGTGTGCTCTTCCGATCTTACCAGGGT-ATCTAATCC (V4R), Taq Phusion (New England Biolabs) and dNTP (New England Biolabs) during 25 cycles (10 s at 98°C, 30s at 45°C, 45s at 72°C). Purity of amplicons was checked on agarose gels before sequencing using Illumina Miseq technology, performed at the Genotoul Get-Plage facility (Toulouse, France). Raw sequences were analyzed using the bioinformatic pipeline FROGS ^21^. Briefly, quality control was performed using Cutadapt (version 1.18) and Flash (version 1.2.11) to delete sequence not between 380 and 500 bp, sequences with ambiguous bases and sequences that did not contain the rightgood primers. Clustering was then performed using Swarm v2.1.2 with an aggregation maximal distance of 3 bases. Chimeras were removed using VSEARCH (v2.9.1) with cross-sample validation. PhiX contamination was removed using VSEARCH. Affiliation was performed using the silva 123 16S database and NCBI Blastn++. A phylogenetic tree was constructed using FastTree (v2.1.10) and sample depth was normalized using GMPR. Physloseq package was used for biostatistical process. The number of observed species and Shannon index (evenness of the species abundance distribution) were calculated to estimate alpha-diversity. Beta-diversity was evaluated by calculating Jaccard distances between samples. Ordination using principal coordinates analyses was performed to represent samples on a 2D plot. Relative abundances of the 5 major phyla were compared within intestinal site using ANOVA. LDA-effect size (LefSe) was then performed to highlight the main differences between IR^fl/fl^ and IR^ΔGUT^ mice caecal microbiota.

### Statistical analyses

Statistical analyses were performed using a non-parametric Mann–Whitney, Two-way ANOVA, T test or χ^2^ analysis for qualitative parameters (Prism10; GraphPad) with a threshold for significance of *P* < 0·05. All data are expressed as mean values with their standard errors.

## RESULTS

### Epithelial IR loss triggers gut leakiness in the absence of obesity and hyperglycemia

To explore the contribution of insulin signaling to intestinal permeability, we generated IR^ΔGUT^ mice with a tamoxifen-inducible deletion of *Insr* in the gut epithelium. We first confirmed the efficiency and specificity of *Insr* deletion. In control IR^fl/fl^ mice, *Insr* mRNA levels in epithelial cells from the upper small intestine (USI) were comparable to liver, but ∼50% lower in the lower small intestine (LSI). By contrast, tamoxifen administration in IR^ΔGUT^ mice markedly reduced *Insr* expression in the gut but not in liver, without major parallel changes in *Igf1r*, confirming both tissue specificity and lack of compensatory mechanisms (**Fig. 1A, Fig. S1A**). At the protein level, IR expression mirrored the mRNA pattern in the control group, whereas nearly absent 10 weeks post-deletion in IR^ΔGUT^ mice. (**Fig. 1B**). Because *Insr* mRNA and IR protein were enriched in the small intestinal crypt compartment compared to the villus in IR^fl/fl^ mice (**Fig. 1C-D**), we further performed *ex vivo* insulin challenge on crypts-derived organoids. While IR^fl/fl^ mice displayed robust AKT phosphorylation, IR^ΔGUT^ mice failed to activate this pathway (**Fig. 1E**), supporting the functional loss of IR-mediated signaling upon gut *Insr* deletion.

**Figure 1:**
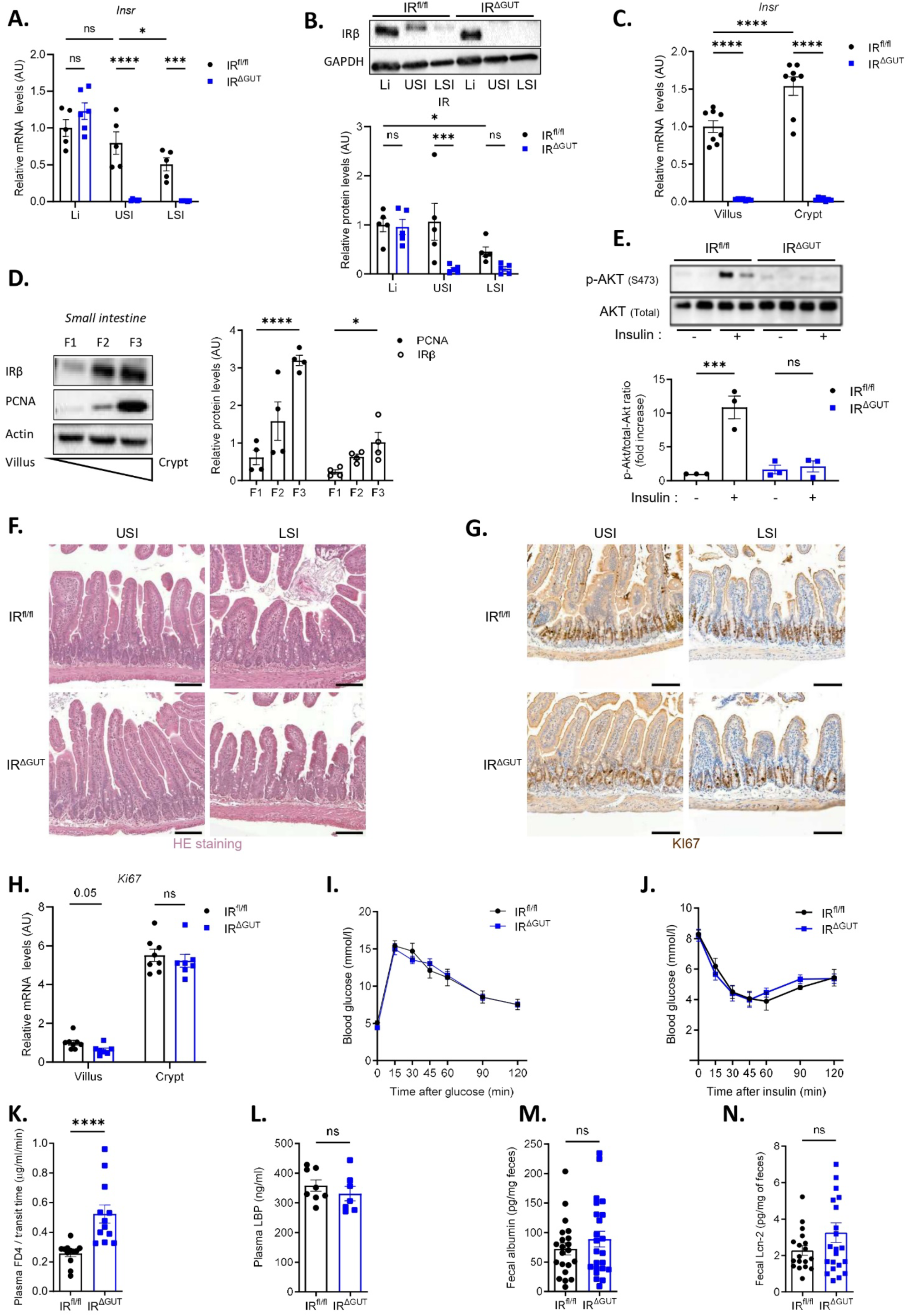
**Intestinal IR loss drives gut leakiness in IR^ΔGUT^ mice**. (A-D) RT-qPCR analysis of *Insr* mRNA levels (A, C) and western blot of IR protein levels by Western blot (B, D). Quantification was performed in epithelial cells isolated from the upper and lower small intestines (USI and LSI, respectively) and the liver (A, B), and along the crypt-villus axis (C, D). (E) Western blot quantification of pAKT/total AKT ratio in intestinal organoids following insulin challenge (100 nM, 5 min). (F-G) Representative images of H&E staining (F), and Ki67 immunostaining of USI and LSI sections (G). Scale bar: 100 µm. (H) RT-qPCR quantification of Ki67 mRNA levels in small intestinal crypts and villi. (I-J) Oral glucose tolerance test (OGTT) and insulin tolerance test (ITT). (K) *In vivo* FD4 assay intestinal permeability assay, (L) plasma LBP levels, (M) fecal albumin concentrations, and (N) fecal lipocalin-2 levels. (A-M) Analyses were performed on both IR^ΔGUT^ and IR^fl/fl^ mice 10 weeks post-tamoxifen treatment. (A, C, H) mRNA levels were normalized to *Gapdh* and *18s* expression. Sample sizes: n = 5–8 (A–L), n = 22-24 (M, N). Statistical analysis: Mann-Whitney test. *p < 0.05; ***p<0,001; ****p < 0.0001.

Given the crucial role of the gut epithelium in glycemic control (via glucose absorption and incretin secretion) and in regulating food intake, we next assessed key metabolic parameters in IR^ΔGUT^ mice. However, no significant differences were observed in body weight, glucose tolerance, or insulin sensitivity upon gut IR deficiency as compared to IR^fl/fl^ control mice (**Fig. 1I-J),** thus allowing the specific role of insulin action to be investigated. Hence, RT-qPCR analyses of epithelial cells isolated from USI and LSI showed comparable mRNA levels of genes encoding either key glucose transporters (*Slc2a2, Slc5a1*), or incretin (*Gcg)* between both genotypes (**Fig. S1F-G, Fig. S1J**). Of note, *Pyy* expression was significantly reduced in the LSI, and *Gip* in the USI of IR^ΔGUT^ mice, consistent with previous reports ^14^ (**Fig. S1H-I**). Moreover, the expression of prototypic hepatic IR-downstream targets (*Fasn*, *Scd1*) remained unchanged following gut IR loss, suggesting that IR signaling regulates a distinct transcription program in this tissue (**Fig. S1D-E**).

Although overall intestinal architecture and crypt proliferation appeared normal in IR^ΔGUT^ mice (**Fig. 1F-H**), FITC-dextran (FD4) assays revealed a significant raise in intestinal permeability, as compared to controls (**Fig. 1K**). Despite epithelial leakiness, detailed analyses using transmission electron microscopy and gene expression profiling showed surprisingly no overt structural defects in tight junction integrity or mucus production by goblet cells (**Fig. S1K-N**). This was also not accompanied by significant change in plasma LBP concentrations, fecal albumin and fecal lipocalin 2 levels, further supporting a non-inflammatory breach in barrier function (**Fig. 1L-N**). As endotoxemia and systemic inflammation are commonly linked to impaired glycemic control in diet-induced obesity, we finally exposed IR^ΔGUT^ mice to a 12-week high-fat diet (HFD) to determine whether such enhanced intestinal permeability could exacerbate inflammatory and metabolic dysfunctions. Although HFD-feeding markedly increased body weight and serum LBP while impairing glucose tolerance, as compared to mice fed a standard chow diet (CD), no significant differences were observed between IR^fl/fl^ controls and IR^ΔGUT^ mice (**Fig. S2A-G**). Taken together, these data demonstrated that gut IR loss is sufficient to enhance epithelial permeability in lean mice, but does not elicit intestinal inflammation or systemic endotoxemia.

### Paneth cell antimicrobial activity is impaired in IR^ΔGUT^ mice

To explore early transcriptional mechanisms underlying gut leakiness in IR^ΔGUT^ mice, we next performed, transcriptomic analyses on intestinal crypts, where IR was enriched (**Fig. 1C-D**), three days after *Insr* deletion. Hence, among the diffentially expressed genes in IR^ΔGUT^ mice (Log2FC < −0.5, p=0.05, 188 and 349 genes significantly down- and up-regulated versus IR^fl/fl^), further analysis of gene ontology (GO) enrichment revealed a marked reduction of terms associated with antimicrobial defense (**Fig. 2A**), a response primarily mediated in the small intestine by Paneth cells. Accordingly, volcano plot analysis showed reduced mRNA levels of several Paneth cell markers (*Lyz1*, *Defa26*, *Mmp7, Ang4, Pla2g2a)* (**Fig. 2B**). This drastic decrease of Paneth cell markers in IR^ΔGUT^ mice was further validated by RT-qPCR (*Lyz1*, *Defa5*, *Defa3*, *Defa6*, *Mmp7)* after both short- and long-term *Insr* deletion (at 3 days and 10 weeks post-tamoxifen, respectively) (**Fig. 2C, Fig. S3C**). Of note, the reduction of lysozyme immunostaining on intestinal section from IR^ΔGUT^ mice was not associated with lower Paneth cells number per crypt, as quantified by immunofluorescence at both time points (**Fig. 2D-F, Fig. S3D-F**).

**Figure 2:**
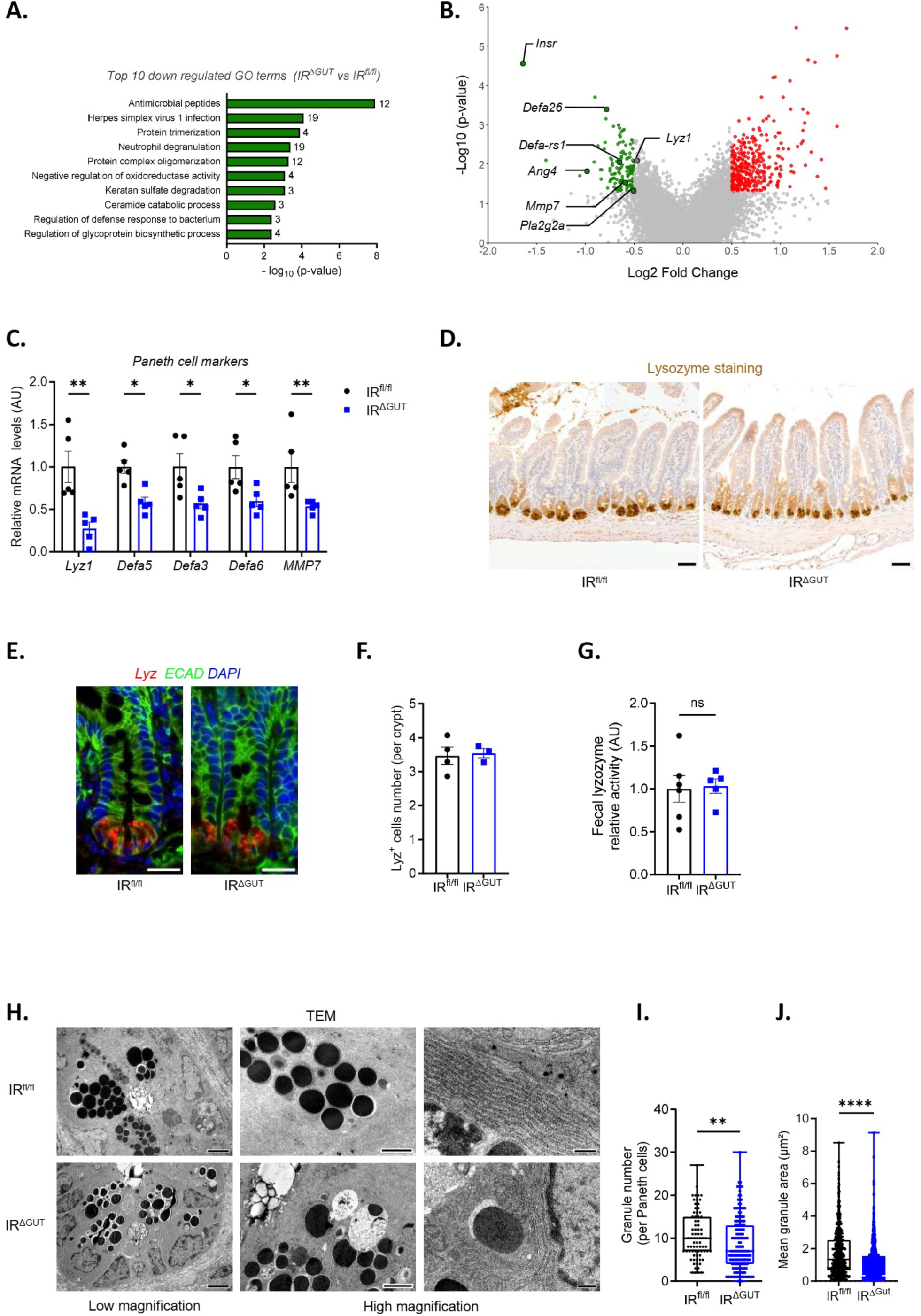
**Paneth cell antimicrobial function is impaired in IR^ΔGUT^ mice**. (A-B) Affymetrix transcriptomic analysis of crypts isolated from IR^ΔGUT^ as compared to their IR^fl/fl^ controls (A) Top 10 downregulated GO terms. (B) Volcano plot of DE genes highlighting significantly downregulated genes associated with Paneth cell identity. (C) RT-qPCR analysis of Paneth cell markers mRNA levels in LSI crypts (as normalized to *Gapdh* and *18s*). (D) Representative images of lysozyme (LYZ) immunostaining on LSI sections. (E) Representative images of immunofluorescence showing ECAD (green) and LYZ (red) staining on DAPI-counterstained (blue) LSI sections. (F) Quantification of Paneth cell number per crypt (10-60 Paneth cells counted per mouse). (G) Fecal lysozyme activity. (H) Representative images of transmission electron microscopy (TEM) analysis on LSI sections. Scale bars: 4 µm (low magnification), 0.5 µm (high magnification). (I) Quantification of Paneth cell granule number and (J) mean granule surface area on TEM images (10-60 Paneth cells analyzed per mouse). (A-J) Analyses were performed in both IR^ΔGUT^ and IR^fl/fl^ mice after 3 days of tamoxifen treatment. Sample sizes: n = 5-6 (A-C, G), n = 3-4 (E, F, H-J). Statistical analysis: Mann-Whitney test. *p < 0.05; **p<0.01 ****p < 0.0001. *p < 0.05; **p<0.01 ****p < 0.0001.

While this suggested that impaired antimicrobial peptides expression results from Paneth cell dysfunction rather than loss, transmission electron microscopy further revealed ultrastructural defects in Paneth cells after both short- and long-term *Insr* deletion. Besides the sustained reduction of Paneth cell secretory granules number and surface area (**Fig. 2H-J, Fig. S3H-J**), early signs of endoplasmic reticulum (ER) stress were observed in Paneth cells, as evidenced by dilated ER membranes shortly after *Insr* deletion (**Fig. 2H**). Indeed, Paneth cells are highly sensitive to perturbations of the ER stress due to their intense secretory activity, and mutations in ER stress-related genes are a risk factor for the breakdown of the intestinal barrier. Given the close interplay between ER stress and autophagy in this cell type, we next evaluated the expression of ER stress (*Bip, Chop, Ero1lb)* or autophagy markers (*Atg5, Atg7, Atg12*) in intestinal crypts isolated from IR^ΔGUT^ mice. However, similar mRNA levels were observed in IR^fl/fl^ and IR^ΔGUT^ mice (**Fig. S3A-B**). Finally, fecal lysozyme activity was measured as an integrated functional readout of Paneth cell antimicrobial capacity, encompassing gene expression, protein production, secretion, and luminal activity. This revealed a significant decrease only after long-term *Insr* deletion (**Fig. 2G, Fig. S3G**), indicating that sustained IR loss is necessary to impair Paneth cell secretory function.

### Paneth cell dysfunction associates with the onset of gut dysbiosis in IR^ΔGUT^ mice

We next performed 16S rDNA sequencing of cecal and fecal microbiota to evaluate whether Paneth cell alterations in IR^ΔGUT^ mice were associated with changes in bacterial composition. Principal component analysis (PCA) revealed that IR^ΔGUT^ mice exhibit a distinct cecal and fecal microbiota profiles compared to control IR^fl/fl^ mice (**Fig. 3A, Fig. S4A**). This shift was associated with an increased number of microbial species in IR^ΔGUT^ mice, with either unchanged or reduced microbial diversity in the cecum and the feces, respectively (**Fig. 3B-C, Fig. S4B-C**). Despite no significant change among major phyla, IR^ΔGUT^ mice exhibited a marked enrichment in *Pseudomonadota* (formerly *Proteobacteria*), a phylum commonly enriched in metabolic or gut inflammatory diseases and associated with deleterious epithelial effects ^20–24^ (**Fig. 3D-F, Fig. S4D**). Moreover, linear discriminant analysis (LDA) of cecal microbiota revealed modest enrichment *Erysipelotrichaceae*, *Erysipelatoclostridiaceae*, *Atopobiaceae* families, alongside increased abundance of *Ca. Stoquefichus*, *Ruminococcus*, *Gordonibacter*, *Oribacteriaceae*, *Turicibacter*, and *Tuzzerella,* and reduced levels of *Ligilactobacillus*, *Anaerotruncus*, and *E. fissicatena* at the genus level (**Fig. 3G**). Feces LDA analysis further confirmed the bacterial family shifts observed in the cecum, while additionally indicating *Monoglobaceae* enrichment and a significant reduction of *Muribaculaceae* (**Fig. S4F**).

**Figure 3:**
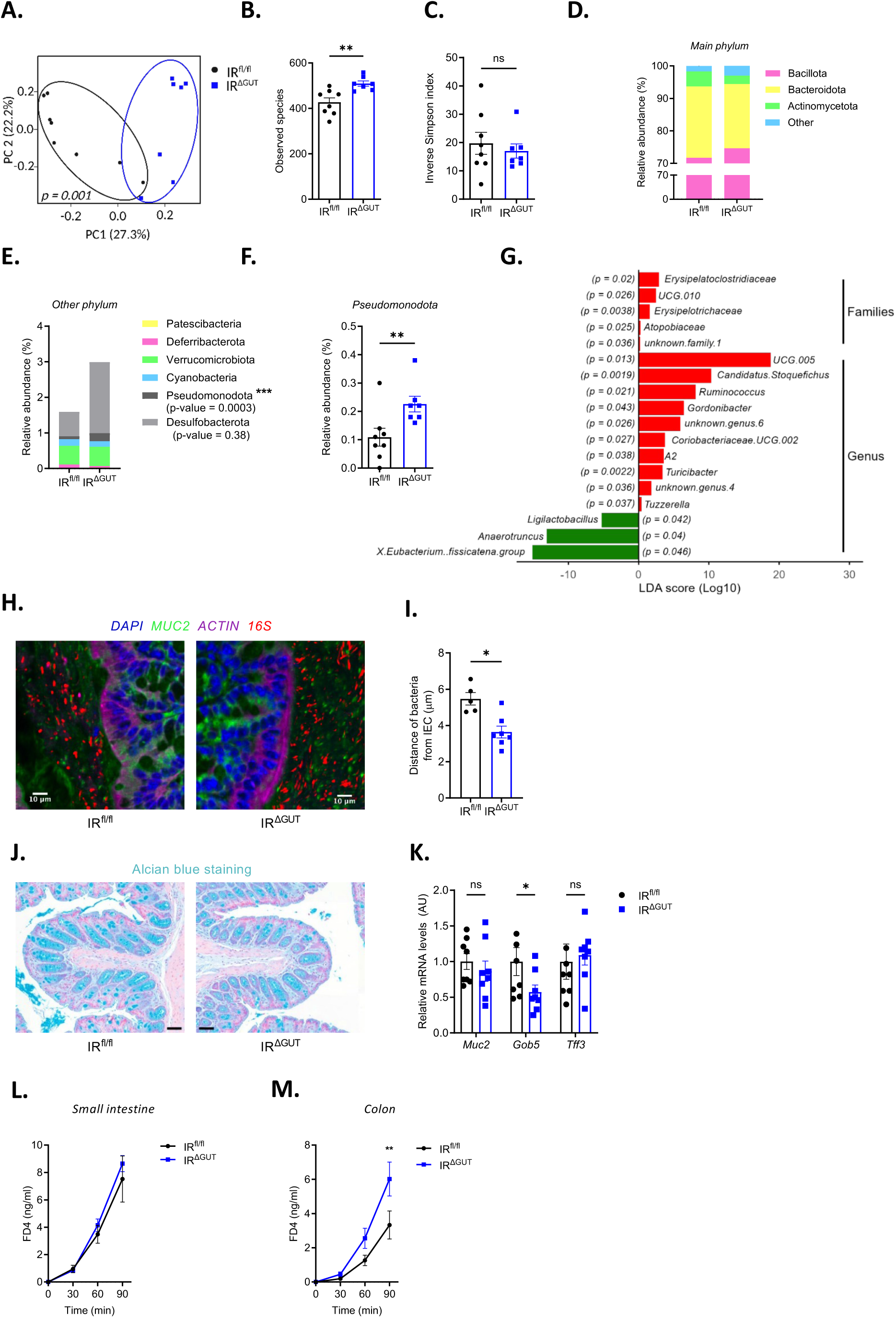
**Gut IR deficiency drives the onset of a distal dysbiosis in IR^ΔGUT^ mice**. (A) Principal coordinate analysis (PCoA) of gut microbiota composition using weighted UniFrac distance matrices, with assessment of (B) bacterial diversity and (C) Inverse Simpson index. (D) Relative abundance of dominant bacterial phyla and (E) expanded view of taxa classified in the “Other” category. (F) Relative m-RNA expression of *Pseudomonadota* abundance in fecal samples. (G) Linear Discriminant Analysis (LDA) scores showing taxa differentially enriched between IR^ΔGUT^ VS IR^fl/fl^. (H) Representative images of bacteria (16S probe *in situ* hybridization, red) embedded in the mucus layer (MUC2 immunostaining, green) overlying the epithelium (ACTIN immunostaining, purple) on DAPI-counterstained colonic sections. (I) Quantification of the distance between the inner mucus layer and the first detectable bacteria (20–30 measurements per mouse). (J) Representative image of Alcian blue staining on colonic sections. (K) RT-qPCR analysis of mucus-related genes in colonic epithelial cells. (L, M) *Ex vivo* FD4 assay permeability assay of small intestine and colon fragments in Ussing chambers. (A-G, L) Analyses were performed on cecal microbiota from IR^ΔGUT^ and IR^fl/fl^ mice 10 weeks post-tamoxifen treatment. Sample sizes: n = 7–8 (A-G, J-L), n = 5–8 (H, I), n = 4–5 (M, N). Statistical analysis: Mann-Whitney test. *p < 0.05; **p < 0.01.

Because besides dysbiosis, increased microbiota encroachment has been reported as both a hallmark and a potential driver of chronic inflammatory disorders, including type 2 diabetes ^25^, we next assessed this feature using 16S rRNA-targeted *in situ* hybridization. This analysis revealed a closer proximity of bacteria to the colonic epithelium of IR^ΔGUT^ mice (**Fig. 3H-I**), without clear evidence of mucolytic bacterial enrichment (data not shown) or overt defects in mucus production (**Fig. 3J-K**). Of note, Ussing chamber analyses revealed that gut leakiness detected *in vivo* in IR^ΔGUT^ mice was confined *ex vivo* to the colon (**Fig.1K**, **Fig. 3L-M**), suggesting that shifts in microbial composition may locally impair colonic epithelial permeability. Finally, since the microbiota is both regulated by and modulates antimicrobial peptides expression ^26,27^, we tested whether defective bactericidal activity of Paneth cells in IR^ΔGUT^ mice was microbiota-dependent. To adress this, IR^fl/fl^ and IR^ΔGUT^ mice were subjected to a broad-spectrum antibiotic treatment prior to tamoxifen administration. Despite nearly complete microbiota depletion in both groups, as measured by 16S rDNA (**Fig. S4H**), *Lyz1* mRNA levels remained significantly lower in IR^ΔGUT^ mice, without changes in mucus-associated genes expression (*Muc2*, *Gob5)* (**Fig. S4I-K**). This suggested that the Paneth cell defect is microbiota-independent, and likely results from a cell-intrinsic effect of insulin signaling loss. Altogether, these findings show that loss of *Insr* in the gut epithelium alone is sufficient to impair Paneth cell function, alterating distal microbiota composition and metabolite profiles, and fostering bacterial encroachment.

### IR loss rapidly alters Lgr5+ ISCs gene signature and dynamics

Consistent with the role of Paneth cells in supporting the intestinal stem cell niche, transcriptomic profiling of intestinal crypts also revealed a significant downregulation of Lgr5^+^ ISC-associated genes (*Lgr5, Olfm4, Soat1, Alcam*) in IR^ΔGUT^ mice 3 days post-tamoxifen administration *in vivo* (**Fig. 4A**). Of note, RT-qPCR analysis further confirmed reduced mRNA levels *Lgr5* and *Olfm4* mRNA levels upon *Insr* deletion (**Fig. 4E**). In line with those observations, OLFM4 immunostaining showed reduced intensity on intestinal section from IR^ΔGUT^ mice (**Fig. 4B**). However, neither the number of OLFM4⁺ ISC per crypt (**Fig. 4C-D**), nor the expression of the proliferation marker Ki67 differed significantly from the IR^fl/fl^ control group (**Fig. S5B-C**). This highlighted that proliferative activity was preserved within IR^ΔGUT^ intestinal crypts despite the downregulation of canonical Lgr5^+^ ISCs markers. Similar analyses after long-term *Insr* deletion *in vivo* showed that Lgr5^+^ ISCs gene signature was restored in IR^ΔGUT^ mice ten weeks post-tamoxifen treatment (**Fig. S5D, S5F**). While this suggested that compensatory mechanisms might underlie the transient reduction of Lgr5^+^ ISCs markers to preserve crypt homeostasis in IR^ΔGUT^ mice *in vivo*, no parallel upregulation of *Hopx* or *Bmi1* mRNA levels, two markers +4 quiescent ISCs, was detected after either short- or long-term *Insr* deletion (**Fig. S5A, S5E**).

**Figure 4:**
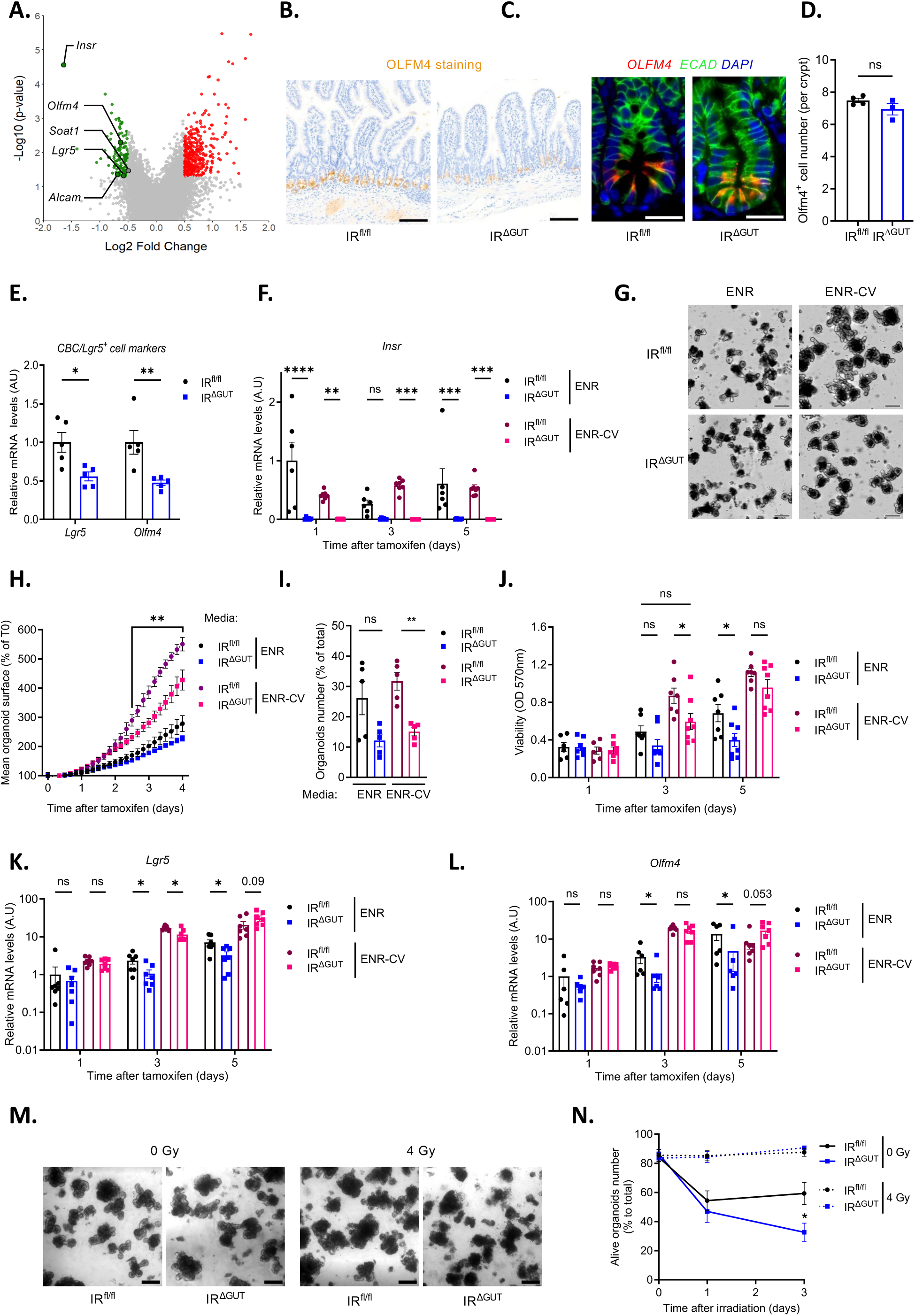
**Gut IR loss impairs Lgr5^+^ ISCs markers and function in IR^ΔGUT^ mice**. (A) Affymetrix transcriptomic analysis of crypts isolated from IR^ΔGUT^ as compared to their IR^fl/fl^ controls. Volcano plot of DE genes highlighting significantly downregulated genes associated with Lgr5^+^ intestinal stem cell (ISC) identity. (B) Representative images of OLFM4 immunostaining on LSI sections. Scale bars: 100µm. (C) Representative images of immunofluorescence showing OLFM4 (green) and LYZ (red) staining on DAPI-counterstained (blue) LSI sections. Scale bar: 20µm. (D) Quantification of ISC number per intestinal crypt (10– 60 ISCs counted per mouse). (E) RT-qPCR analysis of ISC markers mRNA levels in isolated crypts. (F) RT-qPCR analysis of *Insr* mRNA levels in intestinal organoid cultures. (G) Representative images of intestinal organoids at day 4 of culture. Scale bar: 200µm. (H) Quantification of the mean organoid surface area, (I) percentage of large organoids (surface area > 22 980 µm²) among total organoid counts and (J) organoid viability assessed via MTT assay. (K, L) Time-course RT-qPCR analysis of ISC marker mRNA levels during organoid culture. (M) Representative image of intestinal organoid culture 3 days after irradiation. Scale bar: 100µm. (N) Quantification of organoid survival rate. (E-F, K-L) All RT-qPCR results were normalized to *Gapdh* and *18s* expression. (A–E) Analyses were performed on IR^ΔGUT^ and IR^fl/fl^ mice 10 weeks post-tamoxifen treatment *in vivo*. (F–N) Analyses were conducted *in vitro* on IR^ΔGUT^ and IR^fl/fl^ intestinal organoids upon addition of tamoxifen in the culture medium (ENR or ENR-CV) throughout the experimental period (4-5 days). Sample sizes: n = 5 (A–E, N); n = 5– 6 (G–I); n = 6–8 (F, J–L). Statistical analysis: Mann–Whitney test and Two-ways ANOVA (J-L). *p < 0.05; **p < 0.01; ***p < 0.001.

To dynamically assess ISCs defects, we used intestinal organoids, a well-established model of ISCs function ^16^. Organoids were cultured either in ENR medium (EGF, Noggin, R-spondin), which supports physiological epithelial composition and growth, or in ENR-CV medium (ENR supplemented with CHIR99021 and valproic acid), which promotes ISC enrichment ^17^. RT-qPCR analyses confirmed a nearly complete and sustained IR loss *in vitro,* as shown by drastic reduction of *Insr* mRNA levels as early as one day after tamoxifen supplementation in the culture media (**Fig. 4F**). While IR^ΔGUT^-derived organoids exhibited a non-significant trend toward growth defects under ENR conditions, they displayed a progressive reduction in mean surface area and a lower proportion of large organoids under ENR-CV conditions (**Fig. 4G-I, Sup Fig. S5I**), consistent with impaired ISC-driven organoids growth. In line with those observations, expression analysis of ISC markers revealed a significant downregulation of *Lgr5* and *Olfm4* from day 3 in organoids established from IR^ΔGUT^ compared with IR^fl/fl^ controls under both culture media (**Fig. 4K-L**). This was not associated with alteration of either *Hopx* or *Ki67* expression, regardless of the culture conditions (ENR or ENR-CV) (**Fig. S5G**). MTT assays further uncovered reduced organoid viability 5 days post-tamoxifen addition in ENR medium and as early as 3 days in ENR-CV (**Fig. 4J**), suggesting impaired ISCs dynamics in IR^ΔGUT^ mice. Therefore, organoids derived from IR^fl/fl^ and IR^ΔGUT^ mice were next challenged with ionizing radiation to explore the stress-induced regenerative capacity of ISCs. Organoids lacking IR exhibited higher radiosensitivity, as illustrated by a significant increase in apoptotic structures 3 days post-irradiation relative to IR^fl/fl^ controls (**Fig. 4M- N**). Overall, these data show that gut IR loss leads to transient repression of ISC markers without affecting crypt proliferation *in vivo,* while uncovering defective ISC-sustained epithelial growth and reduced survival under stress *in vitro*.

### Paneth cell-derived niche is partially lost in IR^ΔGUT^ mice

Given the disrupted Paneth cell homeostasis previously observed in IR^ΔGUT^ mice (**Fig. 2, Sup Fig. S3**), we next explored whether the altered dynamics of Lgr5^+^ ISCs could stem from a defective Paneth cell-derived niche. After confirming *Insr* mRNA expression in Paneth cells through RNAscope in situ hybridization, following probe validation (**Fig. 5A, Fig. S6A**), we assessed Paneth cell-derived niche factors by RT-qPCR in intestinal crypts isolated from IR^fl/fl^ and IR^ΔGUT^ mice three days and ten weeks after tamoxifen administration *in vivo*. While *Tgfa* and *Dll4* mRNA levels remained comparable between groups, *Wnt3a* and *Egf* showed a trend toward decreased expression after early gut IR loss and returned to control levels by ten weeks, consistent with ISC marker recovery and a potential Paneth cell contribution (**Fig. 5B, Fig. S6B**). To further evaluate niche factors expression specifically in Paneth cells, we first validated their isolation from mouse crypts using a FACS strategy based on established marker combinations (CD45^-^ CD31^-^ Zombie^-^ EPCAM^+^ CD24^+^ ckit^+^ cells) (**Fig. 5C**). The efficient sorting of Paneth cells was confirmed by marked enrichment of *Lyz1*, *Defa6* and *Wnt3a* mRNA levels in the PC^+^ fraction compared to crypt PC^-^ cells (**Fig. 5D**, **Fig. 5F**). Besides revealing higher *Insr* mRNA levels in PC^+^ cells as compared to PC^-^ cells purified from IR^fl/fl^ controls, this approach showed successful deletion of *Insr* in both PC^+^ and PC^-^ fractions isolated from IR^ΔGUT^ mice (**Fig. 5E**). Importantly, *Wnt3a* and *Egf* expression was significantly reduced in Paneth cells (PC^+^) sorted from IR^ΔGUT^ mice relative to the IR^fl/fl^ group (**Fig. 5F, Fig. S6C**), suggesting that IR contributes to niche factors production in this cell type.

**Figure 5:**
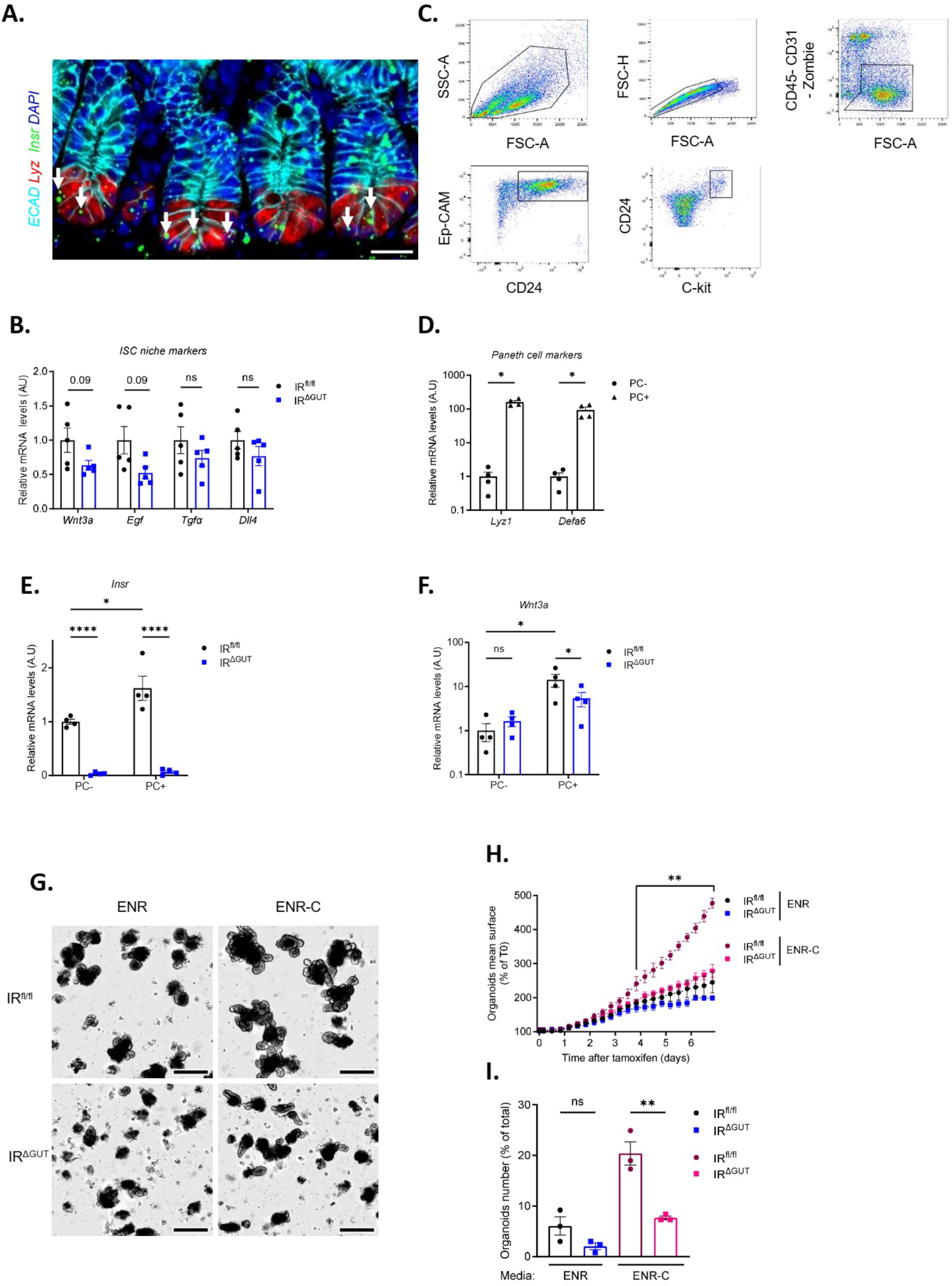
Partial loss of Paneth cell-derived niche in IR^ΔGUT^ mice. (A) RNAscope analysis of *Insr* (green) in Paneth cells (LYZ immunostaining, red) on LSI crypts. Scale bar: 20 µm. (B) RT-qPCR analysis of Paneth cell-derived niche factors in LSI crypts. (C) Flow cytometry gating strategy used to isolate Paneth cells. (D-F) RT-qPCR analysis of Paneth cell niche markers in LSI crypts (D), and of *Insr* (E) and *Wnt3a* (F) mRNA levels in sorted Paneth cells. (G) Representative images of intestinal organoid cultures grown in ENR and ENR-C media. Scale bar: 100 µm. (H) Quantification of mean organoid surface area and (I) percentage of large organoids (surface area > 23,111 µm²) among total organoid counts. (A–F) Analyses were performed in IR^ΔGUT^ and IR^fl/fl^ mice after 3 days of tamoxifen treatment *in vivo*. (G–I) Analyses were conducted *in vitro* on IR^ΔGUT^ and IR^fl/fl^ intestinal organoids upon addition of tamoxifen in the culture medium (ENR or ENR-C) for 7 days. (B-F) RT-qPCR results were normalized to *Gapdh* and *18s* expression. Sample sizes: n = 4–5 per group. Statistical analyses: Mann– Whitney test for panels A–D and H–I; two-ways ANOVA for panels E–F.*p < 0.05; **p < 0.01; ****p < 0.0001.

To test whether these defects underlie the impaired growth of gut organoids derived from IR^ΔGUT^ mice, we supplemented the culture medium with CHIR, a potent Wnt signaling agonist ^28^, without additional EGF since it is already present in standard ENR medium ^16^. In IR^fl/fl^ -derived cultures, mean organoids surface, large organoids proportion as well as *Lgr5* mRNA levels were markedly increased under ENR-C (CHIR99021-supplemented) conditions as compared to ENR (**Fig. 5G-I**, **Fig. S6E-F**). By contrast, IR^ΔGUT^-derived organoids exhibited a limited response to CHIR99021 supplementation, with their overall surface area remaining similar upon ENR and ENR-C conditions throughout the 7-day culture period (**Fig. 5G-I**, **Fig. S6E-F**). Altogether, these findings suggested that the dampened expression of some key niche factors in Paneth cells upon gut IR loss cannot, by itself, fully account for the altered Lgr5^+^ ISCs dynamics observed in IR^ΔGUT^ mice.

### IR sustains homeostasis and metabolic balance in Lgr5⁺ ISCs

Since IR has been identified as part of the Lgr5^+^ ISCs molecular signature ^29^, we next investigated whether gut IR loss induces an intrinsic defect in Lgr5^+^ ISCs homeostasis, independently of the defective Paneth cell-derived niche. After confirming *Insr* expression in Lgr5^+^ ISCs located at the crypt base by RNAscope *in situ* hybridization (**Fig. S7A**), we generated IR^ΔISC^ mice with inducible *Insr* deletion in Lgr5⁺ ISCs by crossing IR^fl/fl^ with *Lgr5-GFP-IRES-CreERT2* mice ^30^. We further employed a FACS strategy based on GFP fluorescence to isolate Lgr5^+^ ISCs (GFP^+^) from other epithelial cells of intestinal crypts (GFP^-^), as validated by the marked enrichment in *GFP* and *Lgr5* mRNA levels in GFP⁺ versus GFP⁻ fractions (**Fig. 6A-B**). Efficient IR loss in Lgr5^+^ ISCs from IR^ΔISC^ mice was demonstrated after three days of tamoxifen administration, as shown by the pronounced reduction of *Insr* mRNA levels in GFP^+^ cells relative to *Lgr5-GFP-IRES-CreERT2* control mice (Ctrl mice) (**Fig. 6C-D**).

**Figure 6:**
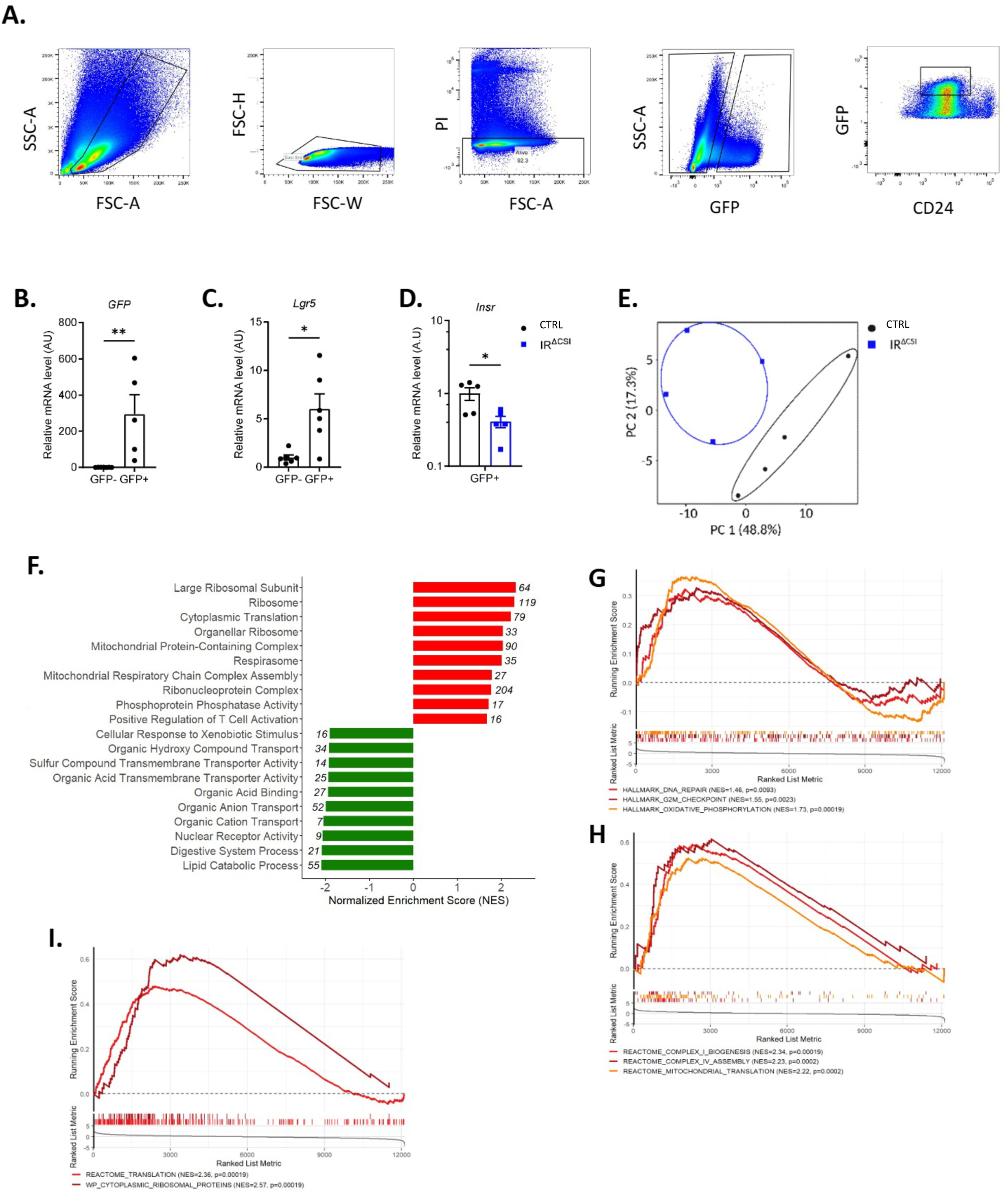
IR deficiency in Lgr5^+^ ISC drives transcriptional reprogramming in IR^ΔISC^ mice. (A) Flow cytometry gating strategy for isolation of Lgr5^+^ intestinal stem cells (ISCs) from crypts, based on GFP expression (GFP^+^ fraction, as compared to GFP^-^ cells). (B, C) Relative mRNA expression of *GFP* (B) and *Lgr5* (C) in GFP^+^ and GFP^-^ fractions of IR^fl/fl^ mice. (D) RT-qPCR analysis of *Insr* mRNA levels in GFP^+^ ISCs. (E) Principal coordinate analysis (PCoA) of transcriptomic profiles from sorted GFP^+^ ISCs. (F) Gene Ontology (GO) enrichment analysis showing the top 10 up- and down-regulated pathways in IR^ΔCSI^ vs IR^fl/fl^ ISC. (G-I) Gene Set Enrichment Analysis (GSEA) illustrating significant upregulation of gene sets involved in DNA repair, G2/M checkpoint control, oxidative phosphorylation (G), mitochondrial complex I and IV biogenesis, mitochondrial translation (H), as well as cytoplasmic translation and ribosomal protein expression (I). (A-I) Analyses were performed in IR^ΔISC^ and/or IR^fl/fl^ mice after 3 days of tamoxifen treatment *in vivo*. Sample sizes: n = 4-5 per group (A-I). Statistical analyses: Mann– Whitney test. *p < 0.05; **p < 0.01.

To explore early transcriptional mechanisms underlying ISCs defects previously observed in IR^ΔGUT^ mice, we next performed comparative bulk RNA sequencing on sorted Lgr5^+^ ISCs (GFP^+^) from IR^ΔISC^ and Ctrl mice. Principal component analysis (PCA) revealed genotype-specific transcriptional signatures, reflected by their distinct clustering (**Fig. 6E**). Gene ontology (GO) enrichment analysis of most upregulated genes in Lgr5+ ISCs isolated from IR^ΔISC^ indicated a strong induction of overall ribosome biogenesis and protein translation, together with mitochondrial metabolic processes including respiratory chain activity (**Fig. 6F**). By contrast, downregulated GO terms were predominantly related to transport and metabolism of organic compounds together with lipids metabolism, in particular sphingolipids and glycero-phospholipids catabolism. These findings were confirmed by Gene Set Enrichment Analysis (GSEA), which highlighted enrichment of multiple metabolic and biosynthetic programs in Lgr5^+^ ISCs isolated from IR^ΔISC^ mice as compared to Ctrl. Upregulated pathways included oxidative phosphorylation, mitochondrial complex I and IV biogenesis, mitochondrial and cytoplasmic ribosomal activity, DNA repair, and G2/M checkpoint regulation (**Fig. 6G-I**). Downregulated pathways included fatty acid metabolism, glycolysis, and glycerosphingolipid catabolism (**Fig. S7B-C**), suggesting reduced metabolic flexibility. Overall, these data demonstrated that IR expression in Lgr5⁺ IScs is critical for sustaining homeostasis and metabolic balance, as its loss induces a transcriptional reprogramming characterized by increased mitochondrial activity and protein synthesis, coupled with reduced lipid metabolism, likely as an adaptive mechanism to support short-term survival.

### Loss of intestinal IR enhances susceptibility to inflammation and enteric infections

Lastly, since IR^ΔGUT^ mice exhibited gut leakiness and impaired barrier function without developing local inflammation under baseline conditions (**Fig. 1K-N**), we next evaluated their susceptibility to either chemical or infectious inflammatory challenges. To this end, IR^ΔGUT^ and IR^fl/fl^ mice were first subjected to dextran sulfate sodium (DSS)-induced epithelial injury and colitis. While body weight remained stable and comparable between IR^fl/fl^ and IR^ΔGUT^ mice during the first 4 days of DSS exposure (day 0 to 4 after DSS administration), IR^ΔGUT^ mice experienced a significant greater weight loss during the 3 days (day 6 to 8) following DSS withdrawal, suggesting poor regenerative capacity (**Fig. 7A**). Compared to controls, IR^ΔGUT^ mice exhibited at day 8 severe epithelial damage and persistent mucosal inflammation, as demonstrated by H&E staining and histological scoring of colonic sections (**Fig. 7B-C**). At day 8, IR^ΔGUT^ mice also showed exacerbated local inflammation, as reflected by increased fecal lipocalin-2 (**Fig. 7D**), elevated colonic MPO activity and pro-inflammatory cytokines (*Tnfα*, *Il6*, *Il17a)* mRNA levels (**Fig. 7E, Fig. S8I**). To investigate the local immune response in IR^ΔGUT^ mice, flow cytometry analyses were performed on immune cell populations isolated from the colonic *lamina propria*. The overall frequencies of neutrophil, monocyte, B cell, T cell or CD45⁺ populations remained comparable between genotypes at day 8 (**Fig. S8A-F**). However, both total and MHC class II-expressing macrophages were significantly reduced in IR^ΔGUT^ mice (**Fig. S8G-H**), suggesting that such impaired antigen presentation and mucosal immune coordination might compromise inflammation resolution.

**Figure 7:**
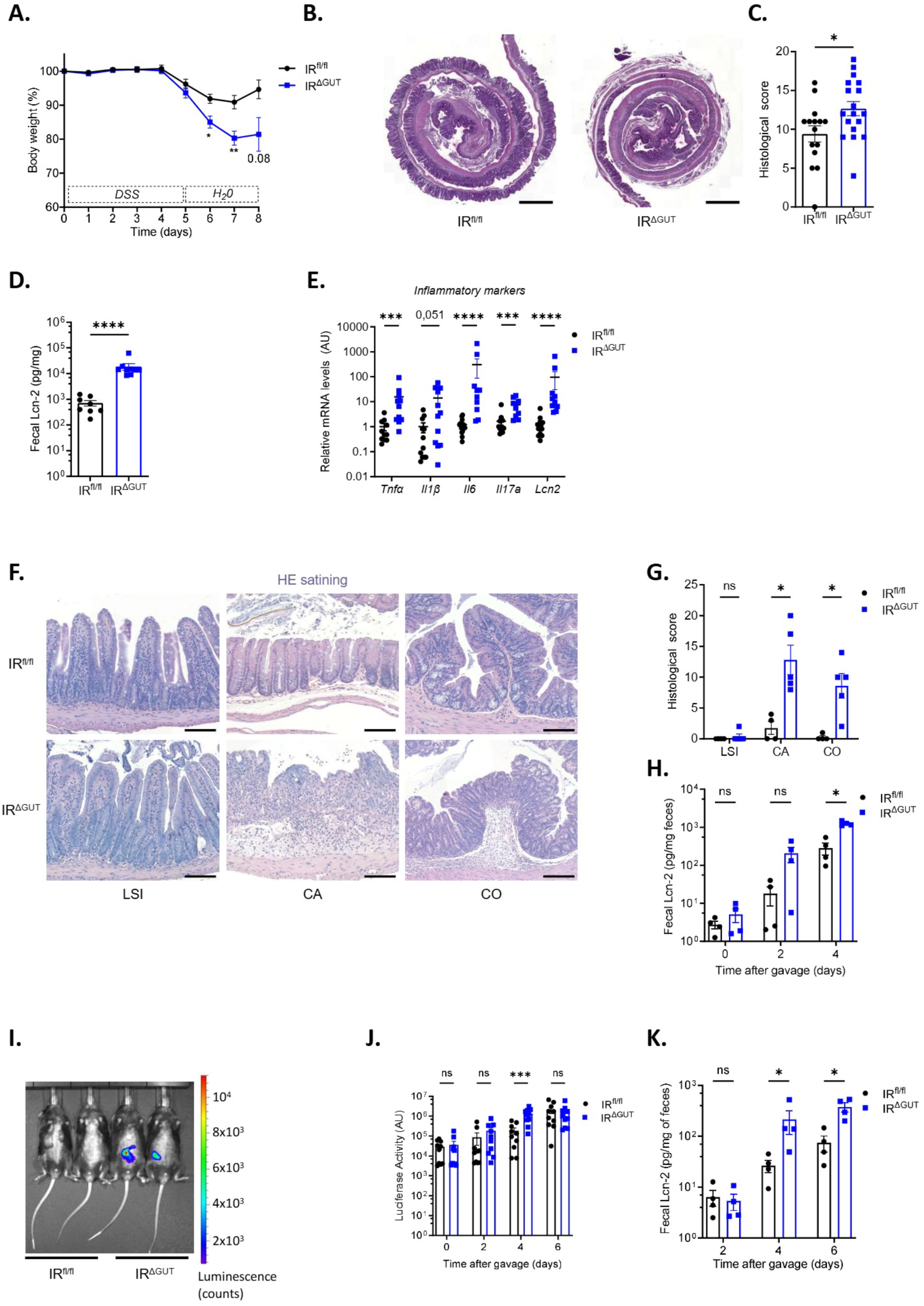
Gut IR loss increases susceptibility to colitis and enteric infections. (A) Percent of body weight loss along the DSS protocol. (B) Representative images of H&E staining of colon rolls at day 8 after DSS treatment initiation. Scale bar: 1mm. (C) Histological scoring of inflammation colon. (D) Fecal lipocalin content in colon at day 8 after DSS treatment initiation. (E) RT-qPCR quantification of inflammatory markers in colon at day 8 after DSS treatment initiation (as normalized to *Gapdh* expression). (F, G) Representative images of H&E staining on LSI, caecum (CA) and colon (CO) epithelium following *Salmonella typhymurium* infection. Scale bar: 100µm. (G) Histological scoring of inflammation in LSI, caecum (CA) and colon (CO). (H) Kinetics of fecal lipocalin (Lnc-2) content after *S. typhymurium* gavage. (I) Representative images of live imaging of *Citrobacter rodentium* colonization 4 days after bacterial infection. (J) Kinetic quantification of *C. rodentium* colonization through live imaging along the infection time. (K) Kinetics of fecal lipocalin content during *C rodentium* infection. (A-K) Analyses were performed in both IR^ΔGUT^ and IR^fl/fl^ mice 10 weeks post-tamoxifen treatment. Sample sizes: n =8-10 (A, D, J), n=14-18 (B, E), n=4-5 (G, H, K). Statistical analysis: Mann– Whitney test. *p < 0.05; ***p<0.001 ****p < 0.0001.

While those findings indicated impaired epithelial regenerative in IR^ΔGUT^ mice, we next evaluated their susceptibility to infection by *S. typhimurium*, a pathogen also triggering strong epithelial damage and inflammation ^31,32^. Colonization efficiency was confirmed by the progressive increase of fecal lipocalin-2 over the course of infection (day 0 to 4 after *S. typhimurium* gavage) in both IR^fl/fl^ and IR^ΔGUT^ groups (**Fig. 7H**). Nonetheless, by day 4, IR^ΔGUT^ mice displayed an exacerbated inflammation compared to controls. This was evidenced by higher fecal lipocalin-2 levels, increased colonic MPO activity, worsened cecal and colonic histological scores, and a non-significant trend toward elevated cytokine expression (*Il6, Il17a*) (**Fig. 7F-H, Fig. S8N-O**). Of note, no significant differences were detected in body weight loss, caecum weight, or colon length during infection (**Fig. S8J-L**). To further extend these findings, we tested the susceptibility of IR^ΔGUT^ mice to *C. rodentium*, a murine enteropathogen that triggers colitis though epithelium adhesion ^33^. Using a bioluminescent *C. rodentium* strain expressing luciferase, we monitored its *in vivo* colonization kinetics. By day 4 post-infection, IR^ΔGUT^ mice exhibited significantly higher luminescence signals, indicative of greater bacterial burden (**Fig. 7I-J**). In line with this, fecal lipocalin-2 levels were also elevated at days 4 and 6 post-infection, supporting increased local inflammation (**Fig. 7K**). Taken together, these results demonstrate that gut IR deficiency heightens susceptibility to colonic inflammation, underscoring a critical protective role of IR signaling in preserving intestinal homeostasis under stress conditions.

## DISCUSSION

While gut permeability defects associated with insulin resistance and hyperglycemia have been shown to arise independently of obesity ^9^, this study further demonstrates that disruption of IR-mediated signaling in the intestinal epithelium, as observed during the diabesity cascade ^11,12^, is sufficient to trigger disrupt selective epithelial permeability. Beyond highlighting IR as a crucial gatekeeper of the selective gut permeability, we identify two major components of the intestinal barrier - Paneth cells and intestinal stem cells (ISCs) - as novel targets of insulin action. Finally, we provide evidence that gut IR loss enhances susceptibility to local inflammation and bacterial infection.

Paneth cells efficiently contribute to intestinal innate immunity, protection against enteric pathogens and bacterial translocation ^34^. Here, we show that IR is expressed in Paneth cells and its inducible deletion in the gut epithelium induces a rapid and prolonged alteration of their homeostasis, as characterized by morphologic alterations of their secretory granules and drastic impairment of their bactericidal capacities. Consistent with this observation, defective gut antimicrobial function has been previously reported in hyperglycemic mice models, including STZ-induced insulin deficiency ^35^ and HFD-induced obesity ^36^, as well as, albeit modestly, upon constitutive deletion of intestinal IR in mouse ^15,37^. Interestingly, Paneth cells are known to respond to local inflammatory cues by acquiring stem-like properties during DSS-induced enteritis ^38^ and by undergoing marked expansion upon *S. Typhimurium* infection ^39^. Consistent Paneth cell defects upon gut IR loss, IR^ΔGUT^ mice parallelly exhibited higher susceptibility to both DSS-induced inflammation and *Enterobacteriaceae* infection, including *S. Typhimurium.* Our data also reveal a causal link between Paneth cell dysfunction and the onset of microbial imbalance in the distal gut of IR^ΔGUT^ mice, underscoring the essential role of IR in maintaining local eubiosis. In line with prior associations in metabolic syndrome, gut IR loss drives a marked expansion of pro-inflammatory *Pseudomonadota,* together with reduction of metabolically beneficial *Bacteroidetes* and *Actinobacteria* species ^4,40,41^. Of note, Ussing chamber analyses revealed that gut leakiness detected *in vivo* in IR^ΔGUT^ mice was *ex vivo* restricted to the colonic epithelium, thereby suggesting that distal dysbiosis following gut IR loss could be at play in this feature. Among potential downstream targets of IR, impaired Paneth cells homeostasis in IR^ΔGUT^ mice may stem from attenuated AKT signaling and subsequent activation of FOXO1 transcriptional activity. Indeed, downregulation of intestinal FOXO1/3 has been shown to promote Paneth cells differentiation both *in vitro* and *in vivo* ^42^. Moreover, given their intense secretory activity, Paneth cells are especially vulnerable to ER stress and autophagy in response to stress ^43,44^, both pathways known to be regulated by insulin signaling ^45^. Importantly, their functional interplay in Paneth cells contribute to the pathogenesis of inflammatory bowel disease ^46,47^. Therefore, beyond the ultrastructural signs of ER stress detected in Paneth cells from IR^ΔGUT^ mice, further investigations are warranted to determine whether defective autophagy and/or ER stress underlie their abnormal secretory phenotype.

A primary means of gut protection against the microbiota is the multi-layered mucus barrier covering the intestinal surface, which keeps the vast majority of bacteria at a safe distance from the epithelium. Interestingly, beyond cecal dysbiosis and associated microbial metabolome shifts, IR^ΔGUT^ mice exhibited higher colonic bacterial encroachment, as previously reported in HFD-induced obese mice and correlated with body mass index and impaired glycemic control in human ^25,48^. Aside from reduced colonic expression of *Gob5*, which deficiency does not significantly compromise mucus barrier or promote bacterial translocation in mice ^49^, no clear evidence supported a significant disruption of mucus production by goblet cells in IR^ΔGUT^ mice. While another study showed a significant alteration of the mucus layer in the colon of STZ-induced hyperglycemic mice - a defect that was reversible by exogenous insulin administration - this model could not disentangle the effects of hyperglycemia from those of impaired epithelial insulin signaling ^50^. According to our data, a proteomic analysis performed on mice carrying a constitutive intestinal deletion of *insr* revealed only minor alterations in colonic Muc2 levels ^13^. Finally, our 16S rRNA sequencing analysis indicated no enrichment of commensal mucolytic bacteria (data not shown), such as *A. muciniphila,* suggesting that enhanced bacterial encroachment observed in IR^ΔGUT^ mice arises from alternative mechanisms. Because the mucus layer is composed of mucins, which glycosylation patterns tightly regulates its physicochemical properties (viscosity, permeability) ^51^, further analyses of mucus rheology and glycosylation may help clarifying whether IR^ΔGUT^ mice exhibit functional alterations of the colonic mucus barrier.

As previously reported upon constitutive gut IR deficienfy ^14,15^, we show that IR loss in the adult mouse gut mucosa did not alter *in vivo* epithelial growth and overall morphology under steady state conditions. However, our data pinpoint that intestinal IR loss impairs both ISCs homeostasis and the Paneth cell-derived niche, as evidenced by decreased expression of ISCs markers (*Lgr5*, *Olfm4),* impaired growth and survival of intestinal organoids, as well as reduced expression of niche factors (*Wnt3a*, *Egf).* Besides bactericidal defenses, Paneth cells are indeed critical for creating an antimicrobial and nurturing environment that sustains ISCs renewal and gut epithelial integrity. In this context, supplementing the culture media with a pharmacological Wnt activator to restore defective Paneth cell-derived niche ^16,52,53^ failed to rescue the growth defects of gut organoids derived from IR^ΔGUT^ mice. may affect either a combination of Paneth cell–derived niche factors or ISCs directly, independently of the niche. Of note, Paneth cells were shown to primarily rely on glycolysis - a metabolic pathway commonly controlled by insulin - thereby delivering lactate to adjacent ISCs, where it fuels oxidative phosphorylation ^54^. Yilmaz et al. also proposed that mTORC1 - an IR downstream target - acts as a nutrient sensor in Paneth cells thereby enabling the adaptation of ISCs function to nutritional status ^55^. Hence, impaired epithelial insulin signaling may compromise the metabolic interaction between Paneth cells and ISCs. However, the hypothesis of a direct insulin action in ISCs is supported by earlier studies identifying IR and its substrate IRS1 as part of the molecular signature of Lgr5⁺ ISCs ^29^. In line with this, our transcriptomic data on IR^ΔISC^ mice indicate that IR deficiency specifically in Lgr5*^+^* ISCs rapidly led to a transcriptional reprogramming toward enhanced mitochondrial respiratory activity and protein synthesis, marked by upreguled oxidative phosphorylation (OXPHOS), mitochondrial biogenesis, and ribosomal pathways. Given the established reliance of Lgr5*^+^* ISCs identity on both ribosome dymanics ^56,57^ and OXPHOS ^54,58^, this suggested that the shift of IR-deleted Lgr5*^+^* ISCs towards an OXPHOS-dependent state might represent a compensatory survival mechanism. Moreover, IR loss in Lgr5*^+^* ISCs was accompanied by an early downregulation of glycolysis and lipid catabolism (notably sphingolipids and glycerophospholipids), indicating reduced metabolic flexibility of IR- deleted Lgr5⁺ ISCs. Consistent with this assumption, IR deletion is well known to dampen glycolysis and fatty acid oxidation in the liver, via AKT inactivation and FOXO1/3 nuclear retention. This phenomenon increases reliance on amino acid catabolism as an energetic substrate and is characterized by elevated expression of *Klf15* ^59^. Notably, our RNA-seq data show a strong increased *Klf15* expression (data not shown), indicating that IR-deficient ISCs may shift towards amino acid metabolism to fuel OXPHOS via NADH and FADH₂ production. Taken together, these findings support a direct role of IR-mediated signaling as a key regulator of ISC homeostasis, likely by safeguarding ISC survival and metabolic fitness under stress conditions.

Interestingly, we show that intestinal IR loss is associated with impaired epithelial regenerative capacities, upon intestinal organoids irradiation *ex vivo* as well as following DSS-induced intestinal inflammation. This regenerative response involves a finely tuned, self-limiting series of cellular and molecular events orchestrated by the transient activation of Lgr5^+^ ISCs and the plasticity of other intestinal epithelial cells. Among the IR-downstream mechanisms that might be at play in defective regenerative response of IR^ΔGUT^ mice, the transcription factor FoxO was shown to regulate the expression of *Dsc3*, which encodes a desmosomal cadherin controlling intestinal integrity in response to inflammatory challenge ^12^. Another interesting candidate is the YAP/TAZ signaling pathway, which acts as a critical gatekeeper of gut epithelial regeneration. Upon DSS or radiation induced-damage, increased YAP activity in regenerating crypts indeed triggers the activation of a transcriptional program driving cell proliferation/survival while preventing over-activation of the Wnt pathway to avoid excessive conversion of Lgr5^+^ ISCs into Paneth cells ^60–62^. Interestingly, while insulin restrains YAP transcriptional activity by promoting its phosphorylation and preventing nuclear translocation ^63^, disruption of DSS-induced YAP-dependent transcriptional program was previoulsy reported in the absence of IR signaling ^64^. Beyond its essential role in sustaining homeostatic Lgr5⁺ ISC proliferation and epithelial turnover, Wnt signaling also orchestrates regenerative response to gut injury ^38,65,66^. Interestingly, IR-mediated inhibition of Wnt signaling is well established as a self-regulatory mechanism in hepatic homeostasis, as downtream AKT activation drives GSK3β inactivation and promotes Wnt ligand secretion via SCD1-dependent palmitoylation ^67^. Thus, more detailed analyses will help delineating the signaling pathways underlying ISC defects upon IR loss.

Overall, this study emphasizes that impaired gut IR signaling ^11^ might contribute to the intestinal barrier dysfunctions associated with the diabesity cascade. We further highlight Paneth cells and intestinal stem cells as previously unrecognized targets of insulin action. By supporting local antimicrobial defenses and epithelial regeneration, we finally demonstrate that intestinal IR acts as a safeguard against local inflammatory and bacterial threats. In this context, clinical evidence support enhanced mucosal infection in diabetic patients ^68–70^. Moreover, insulin intra-rectal instillation reduced inflammation in a pre-clinical mouse model of colon cancer-associated colitis^64^. Therefore, we anticipate that our study will provide promising perspectives for comprehensively mapping the contribution of gut IR signaling to diabetic complications.

## ACKNOWLEGMENTS

We would like to acknowledge Dr Nathalie Rolhion (Centre de Recherche Saint-Antoine, Paris) for providing the *S.Typhymurium* strain and for her fruitful advices on the infection protocol. We are grateful to Béatrice Romagnolo (Institut Cochin, Paris) and Aline Stedman (Institute of Biology Paris-Seine, Paris) for their scientific discussions, and to Gwenaëlle Randuineau (Institut Numecan, Rennes) for her help with microbiota analysis. We also thank the staff of the animal facilities at MOUSET’IC (Institut Cochin) and RPPA (Paris-Sorbonne Université), and the core facilities at the Institut Cochin (HISTIM, IMAG’IC, GENOM’IC, PIME).

Finally, this study was supported by the following grants : European Foundation for the study of Diabetes (EFSD/Novo Nordisk Programme 2014), National Agency for Research (ANR-20-CE14-0044-01, GutBarrIR), Foundation for Medical Research (FRM), French Diabetes Society (SFD/Roche Diabetes Care 2018), University Paris Cité (IdEx Emergence 2020, InidEx DiabetEx 2025, DHU AUTHORS 2016), G.E.N.E graduate school (G.E.N.E. Mentors program 2024), Institut Cochin (Projet inter-équipe 2017).

## SUPPLEMENTAL FIGURES LEGENDS

**Figure S1:**
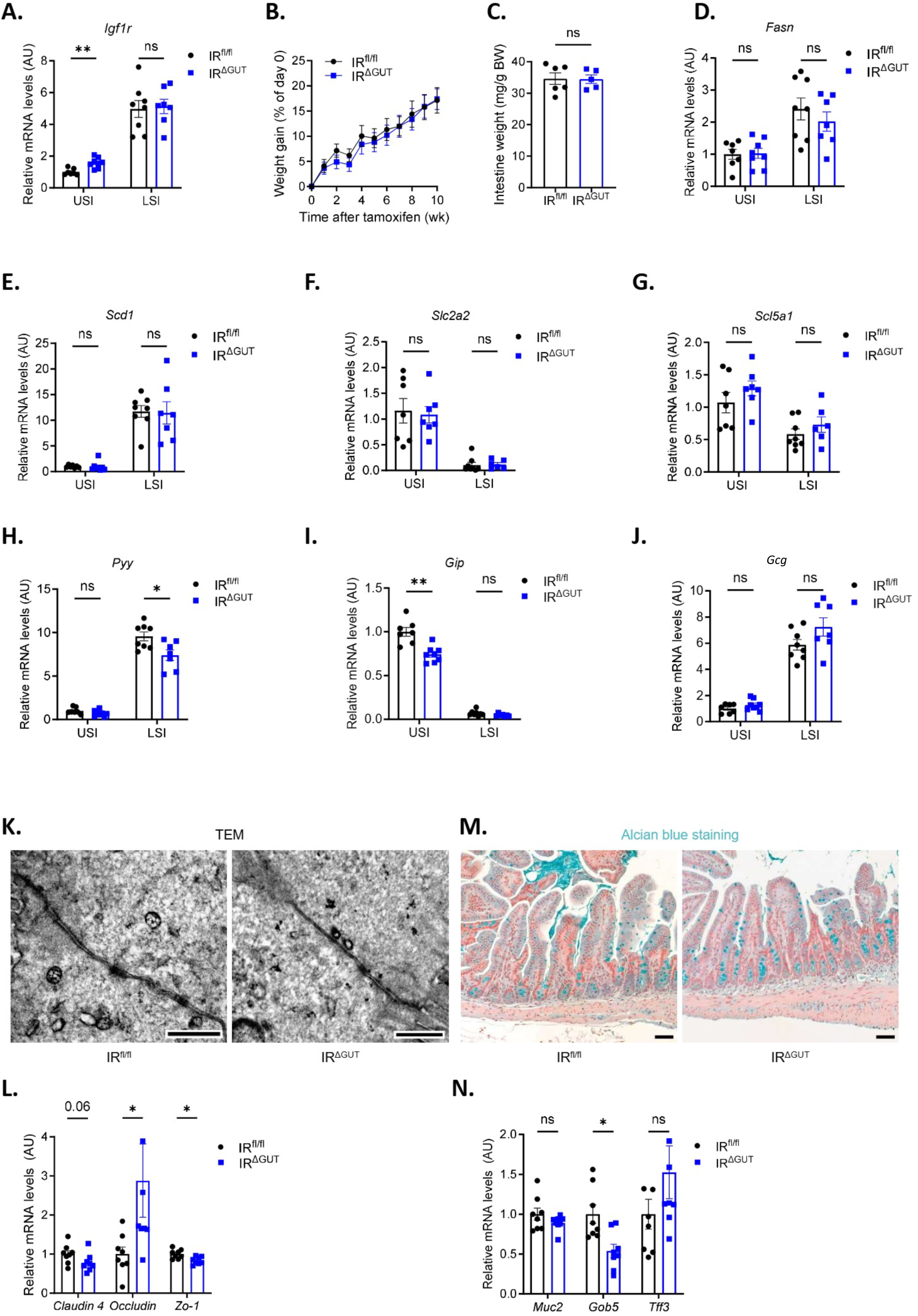
**Absence of gut metabolism and barrier components overt defects in IR^ΔGUT^ mice**. (A) Relative mRNA expression of Igf1r. (B) Body weight gain over 12 weeks following tamoxifen treatment and (C) Intestinal tissue weight at endpoint. Relative mRNA expression of (D-E) lipogenic genes: (D) *Fasn*, (E) *Scd1*, (F-G) glucose transporters (F) *Slc2a2* and (G) *Slc5a1* and (H-J) enteroendocrine hormone transcripts: *Pyy* (H), *Gip* (I) and *Gcg* (J). (K) Transmission electron microscopy (TEM) images of ileal epithelium, scale bar: 0.5 µm. (L) mRNA levels of epithelial junction markers. (M) Representative images of Alcian Blue staining of ileal sections (scale bar: 50 µm) and (N) relative mRNA expression of goblets cells markers. All analyses were conducted in IR^ΔGUT^ and IR^fl/fl^ mice 10 weeks post tamoxifen treatment, except (B). All RT-qPCR results are normalized to *Gapdh* expression and done in isolated IEC (A, D-J, L, N). Sample sizes: n = 5-8 (A, C-N), n=10-12 (B). Statistical analysis: Mann–Whitney test. *p < 0.05; **p<0.01.

**Figure S2:**
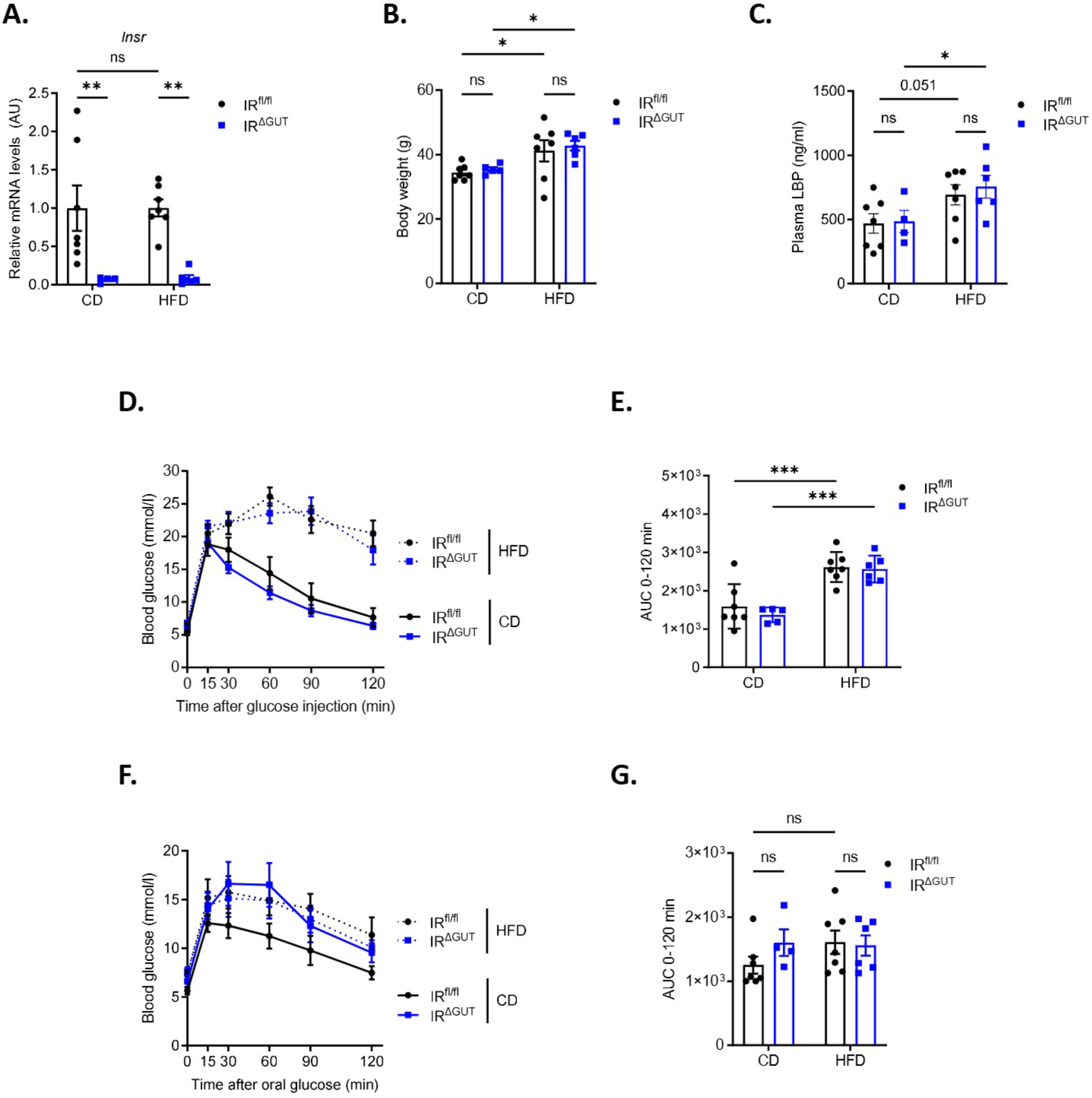
**HFD-induced metabolic alterations are not exacerbated by gut IR loss**. (A) Relative *Insr* mRNA expression in IEC normalized to *Gapdh*. (B) Body weight measurement. (C) Plasma levels of lipopolysaccharide-binding protein (LBP). (D) Blood glucose levels during oral glucose tolerance test (OGTT), with (E) area under the curve (AUC) quantification. (F) Blood glucose during insulin tolerance test (ITT), with (G) corresponding AUC. All analyses were performed in IR^ΔGUT^ and IR^fl/fl^ mice 10 weeks after tamoxifen treatment and after a minimum of 10 weeks under control diet (CD) or high-fat diet (HFD). Sample sizes: n = 4–7 per group. Statistical analysis: Two-way ANOVA. p < 0.05; *p < 0.01; **p < 0.001.

**Figure S3:**
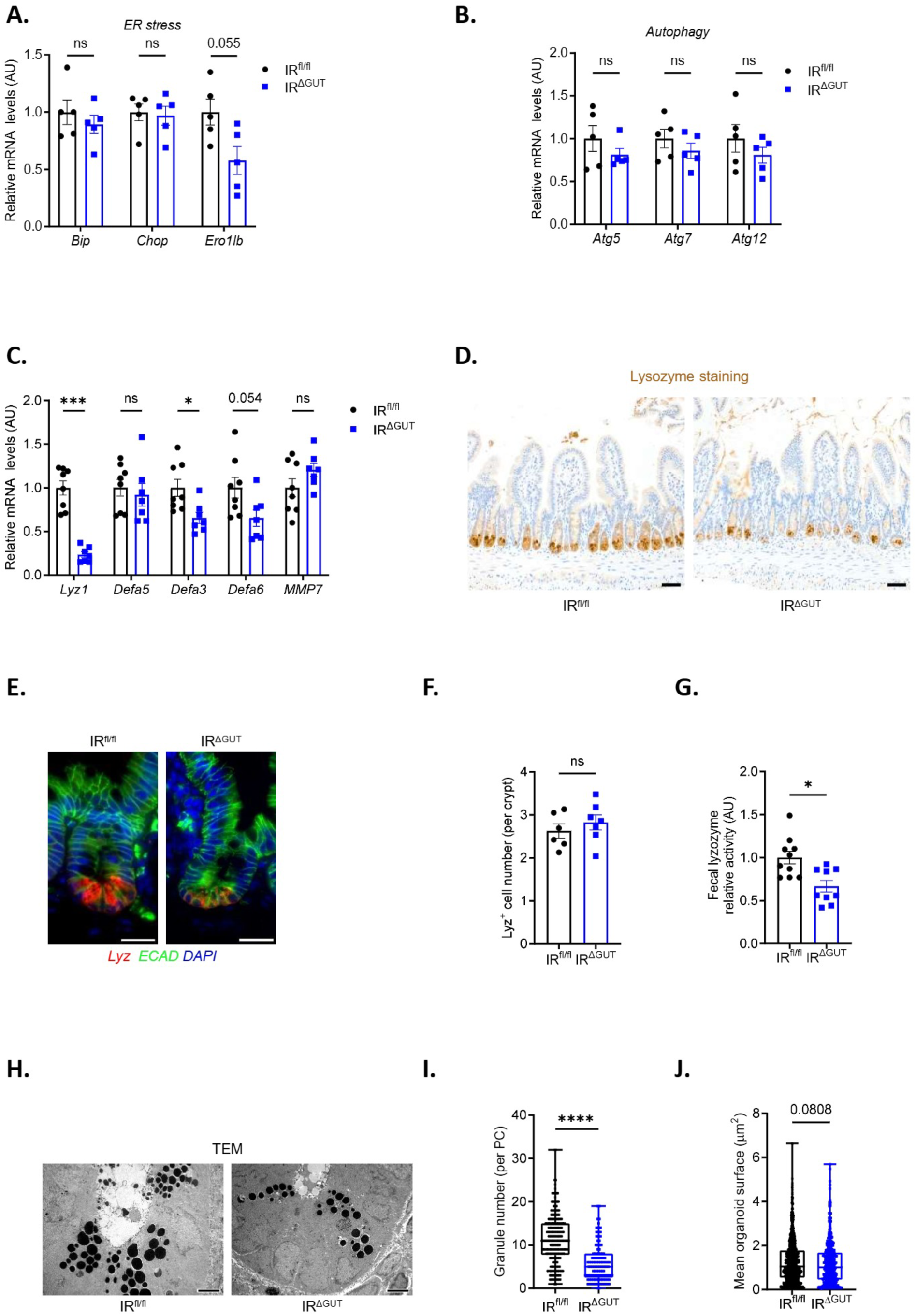
**Impaired Paneth cell function is maintained upon long term gut IR deficiency**. Relative mRNA expression of markers related to (A) endoplasmic reticulum (ER) stress, (B) autophagy, and (C) Paneth cell. Representative images of lysozyme expression in the ileum detected by (D) immunohistochemistry (scale bar: 50 µm) and (E) immunofluorescence (scale bar: 20 µm). (F) Quantification of Paneth cell number per crypt (37–45 Paneth cells counted per mouse). (G) Lysozyme enzymatic activity in fecal samples. (H) Transmission electron microscopy (TEM) images of ileal sections (scale bar: 4 µm). Quantification of (I) Paneth cell granule number and (J) mean granule surface area based on TEM analysis (10–60 Paneth cells analyzed per mouse). All RT-qPCR results (A-C) are normalized to *Gapdh* and *18s* expression and from isolated intestinal LSI crypts. All analyses were performed in IR^ΔGUT^ and IR^fl/fl^ mice. Panels (A–B) were performed after 3 days of tamoxifen treatment; all others were assessed 10 weeks post-treatment. Sample sizes: n = 5 (A–B), n = 3–4 (F), n = 7–8 (C–E, G– J). Statistical analysis: Mann–Whitney test. p < 0.05; **p < 0.001; ***p < 0.0001.

**Figure S4:**
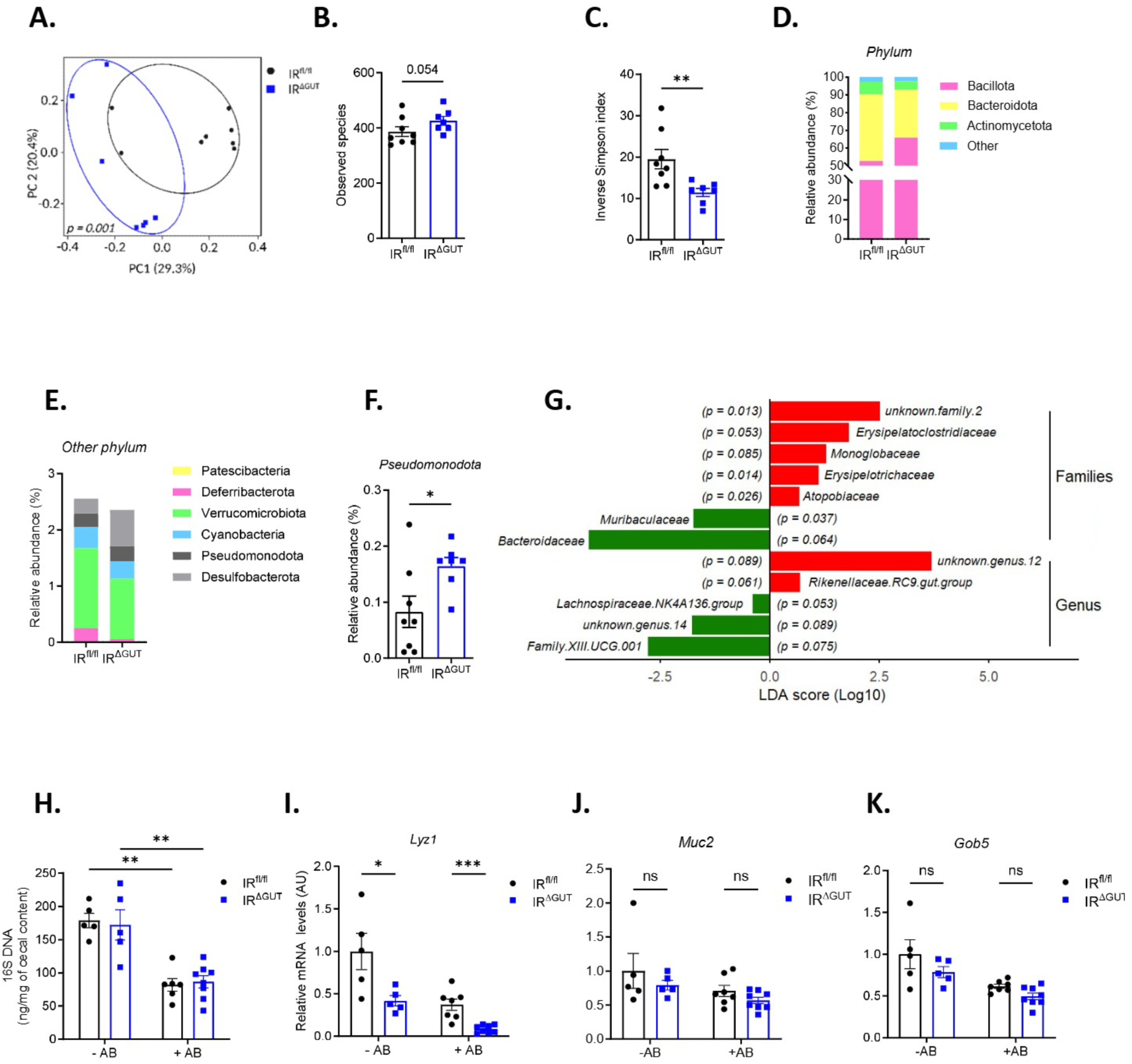
**Intestinal IR loss drives colonic microbial imbalance**. (A) Principal coordinate analysis (PCoA) of colon microbiota composition using weighted UniFrac distance matrices, with assessment of (B) bacterial diversity and (C) Inverse Simpson index. (D) Relative abundance of dominant bacterial phyla and (E) expanded view of taxa classified in the “Other” category. (F) RT-qPCR quantification of *Pseudomonadota* abundance in fecal samples normalized through *Gapdh* expression. (G) Linear Discriminant Analysis (LDA) scores showing taxa differentially enriched between IR^ΔGUT^ VS IR^fl/fl^. (H) Caecal bacterial abundance and relative mRNA expression of (I) *Lyz1*, (J) *Muc2*, and (K) *Gob5* from LSI IEC from mice treated with or without broad-spectrum antibiotics (administered before tamoxifen treatment for a total of 5 days). All analyses were conducted in colon of IR^ΔGUT^ and IR^fl/fl^ mice 10 weeks after tamoxifen treatment, unless otherwise indicated (H–K). Sample sizes: n = 7–8 (A–G), n = 5 (H– K).p < 0.05; *p < 0.01; **p < 0.001. Statistical analyses: Mann–Whitney test (B–F, I–K) and two-way ANOVA (H).

**Figure S5:**
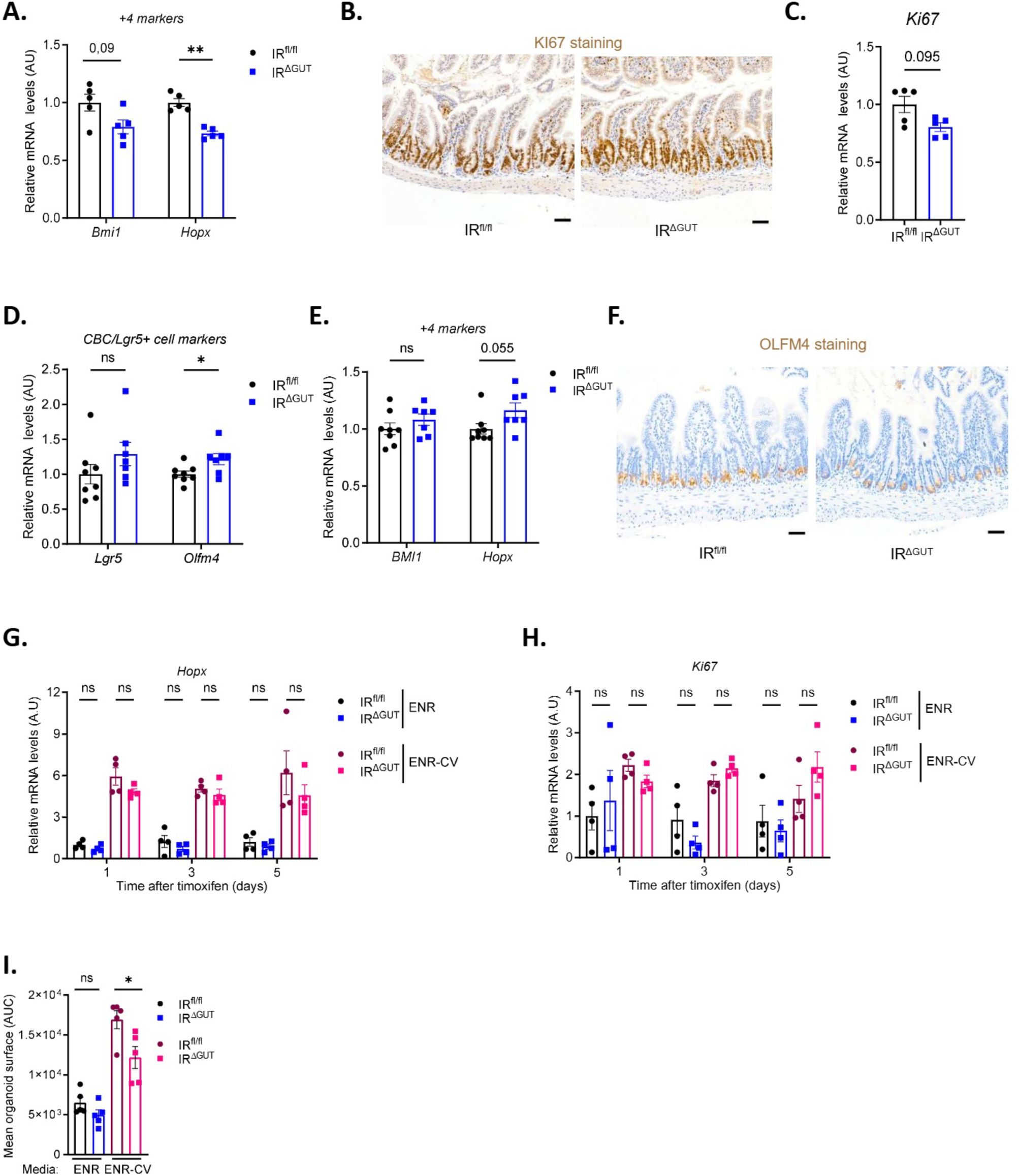
**Absence of compensatory increase of +4 ISC markers in IR^ΔGUT^ mice**. (A) Relative mRNA expression of +4 ISC (C) Ki67 in ileal crypts and (B) Representative image of Ki67 immunohistochemistry in ileal sections (scale bar: 50 µm) 3 days after tamoxifen treatment. Relative mRNA expression of (D) crypt base columnar (CBC) Lgr5⁺ ISC marker and (E) +4 ISC markers 10 weeks post-tamoxifen treatment. (F) Representative immunohistochemistry of *Olfm4* expression in ileal sections, 10 weeks post-treatment (scale bar: 50 µm). Relative mRNA expression of (G) *Hopx* and (H) *Ki67* in intestinal organoids cultured. (I) Area under the curve (AUC) analysis of mean organoid surface area after 5 days of culture. All RT-qPCR results (A, C-E, G-H) are normalized to *Gapdh* and *18s* expression. All analyses were conducted in IR^ΔGUT^ and IR^fl/fl^ mice. *In vitro* experiments (G–I) were conducted in the continuous presence of tamoxifen throughout the experimental period (4–5 days) in ENR or ENR-CV conditions. Sample sizes: n = 4–5 (A–C, G–I), n = 8 (D–F). Statistical analysis: Mann–Whitney test. *p < 0.05; **p < 0.01.

**Figure S6:**
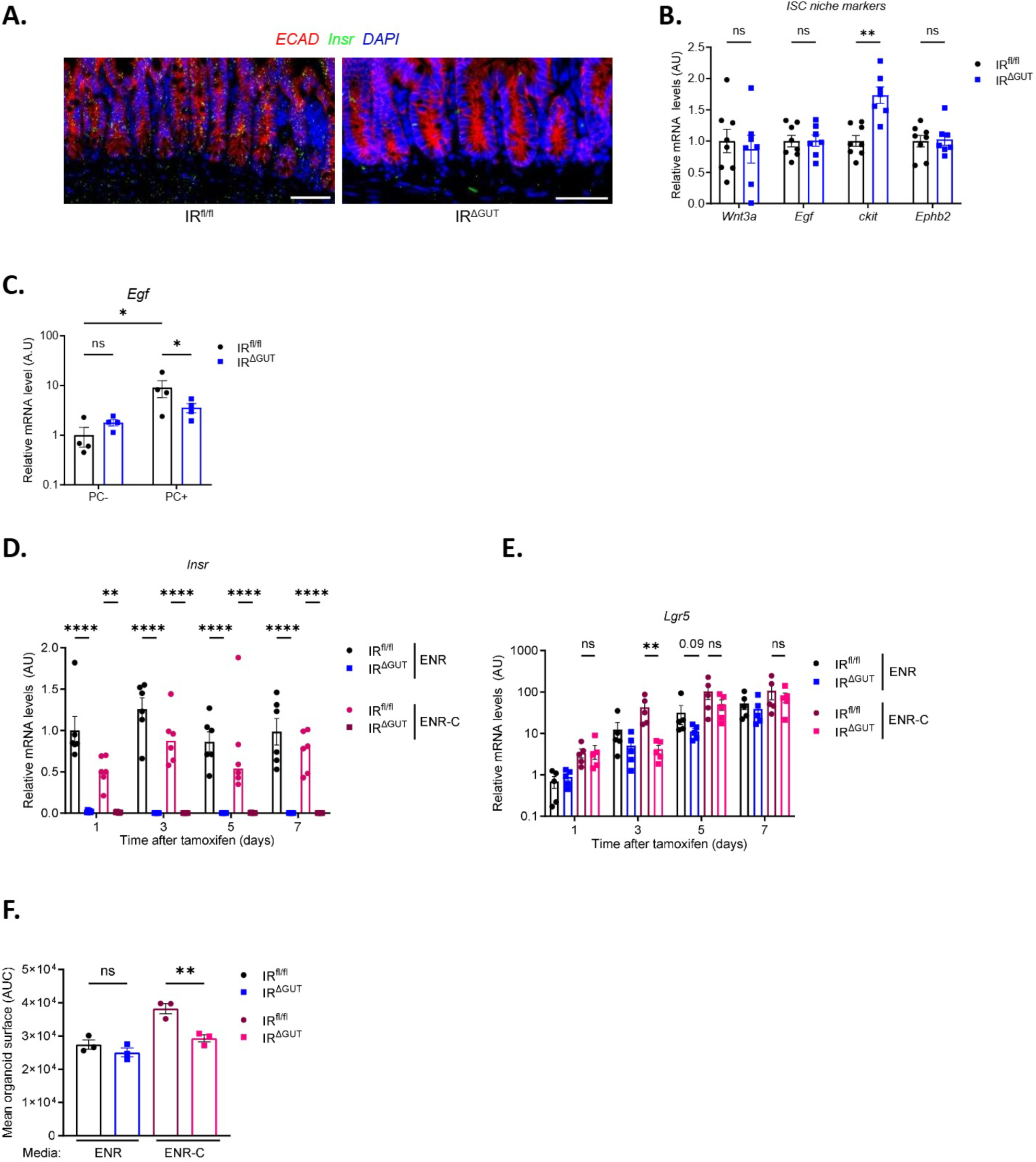
**Lgr5^+^ ISC defects in IR^ΔGUT^ mice are not rescued *in vitro* by epithelial niche supplementation**. (A) RNAscope validation of *Insr* probe (green) in ileal sections, Scale bar: 20 µm. Relative mRNA expression of (B) epithelial niche markers in ileal crypts, (C) Egf in sorted Paneth cells. Relative mRNA expression of (D) *Insr* and (E) Lgr*5* in intestinal organoid cultures. (F) Area under the curve (AUC) analysis of mean organoid surface area after 7 days of culture. All RT-qPCR results (B-E) are normalized to *Gapdh* and *18s* expression. All analyses were performed in both IR^ΔGUT^ and IR^fl/fl^ mice. *In vivo* experiments (A–C) were performed 3 days post tamoxifen treatment. *In vitro* experiments (D–F) were conducted in the continuous presence of tamoxifen in the culture medium for 7 days under ENR and ENR-C conditions. All RT-qPCR data (B–E) were normalized to *Gapdh* and *18s* expression. Sample sizes: n = 7–8 (B, D–E), n = 3–4 (A, F). Statistical analysis: Mann–Whitney test for panels B, D–E; two-way ANOVA for panel C. p < 0.05; *p < 0.01; **p < 0.001; ***p < 0.0001.

**Figure S7:**
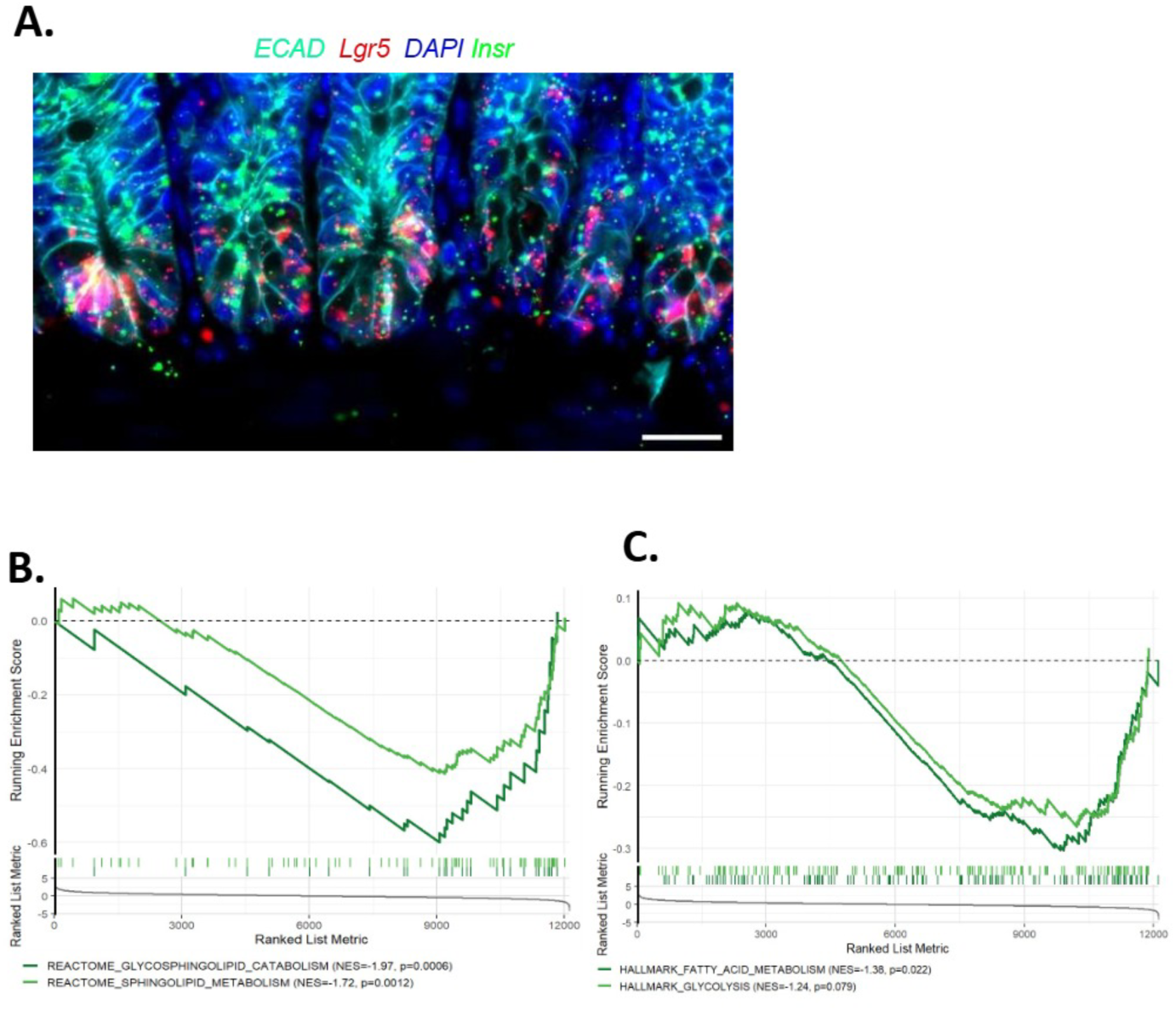
**IR deficiency in Lgr5^+^ ISCs impairs their lipid metabolism**. (A) RNAscope analysis of *Insr* (green) expression in ISC (red) within ileal crypts. Scale bar: 20 µm. Gene Set Enrichment Analysis (GSEA) showing significant downregulation of gene sets involved in (B) glycosphingolipid catabolism and sphingolipid metabolism, and (C) fatty acid metabolism and glycolysis. All analyses were performed in vivo after 3 days of tamoxifen treatment in IR^ΔCSI^ and IR^fl/fl^ mice. Sample sizes: n = 4–5 per group. Statistical analysis: Mann–Whitney test. p < 0.05; **p < 0.001.

**Figure S8:**
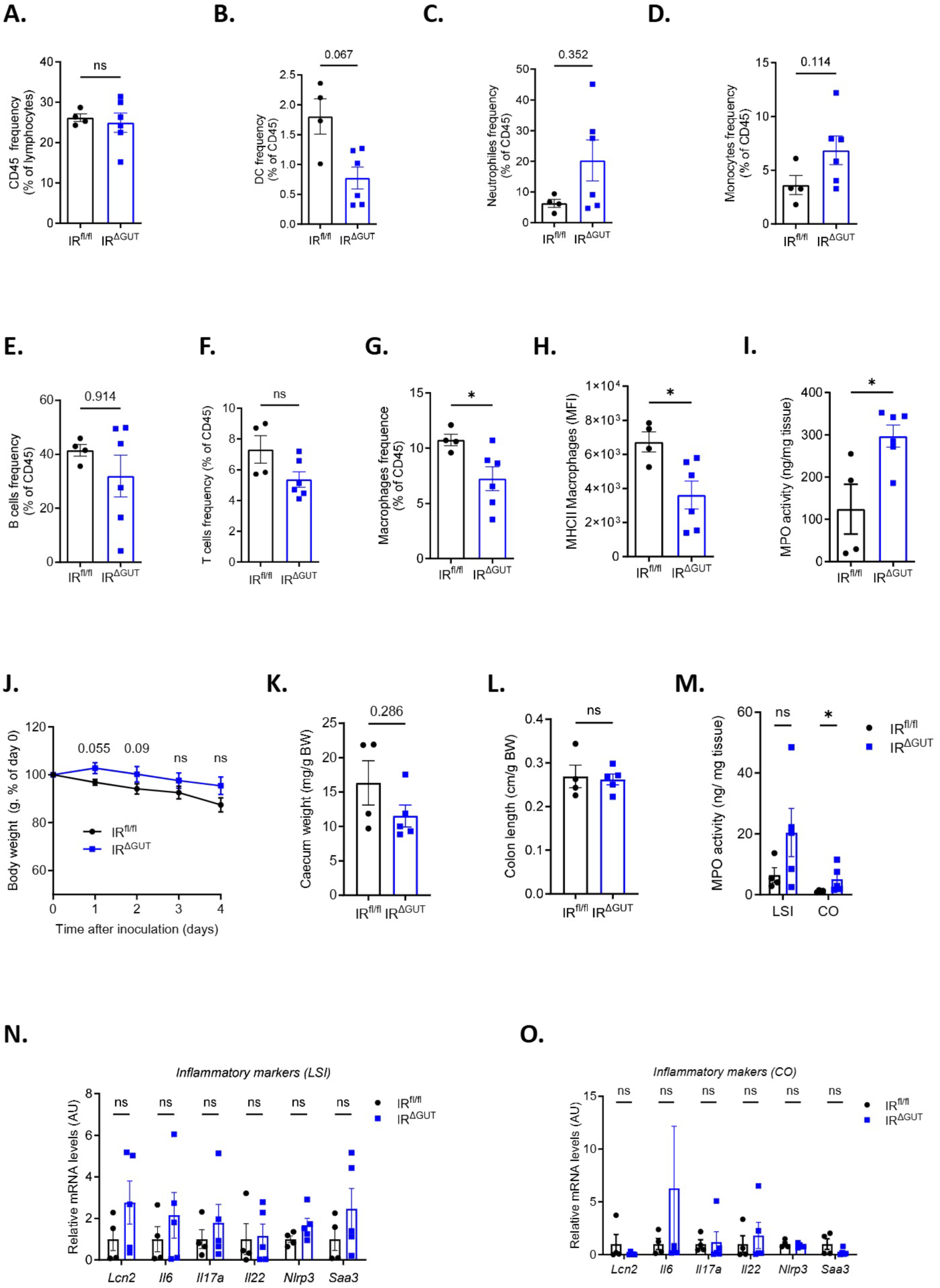
**Gut IR loss increases susceptibility to colitis and enteric infections**. Flow cytometry analysis of immune cell populations within the colonic lamina propria 8 days after DSS administration: (A) frequency of CD45⁺ cells, and proportions of (B) dendritic cells, (C) neutrophils, (D) monocytes, (E) B cells, (F) T cells, and (G) macrophages among CD45⁺ cells. (H) Frequency of MHC class II⁺ CX3CR1⁺ CD11b⁺ F4/80⁺ macrophages. (I) Myeloperoxidase (MPO) activity in colon tissue. In the context of *Salmonella typhimurium* infection: (J) body weight evolution, (K) caecum weight, (L) colon length, (M) MPO activity in the lower small intestine (LSI) and colon, and relative mRNA expression of inflammatory markers in (N) LSI and (O) colon normalized to *Gapdh*. All experiments were conducted *in vivo* in IR^ΔGUT^ and IR^fl/fl^ 10 weeks post tamoxifen treatment. Sample size: n = 4–6 per group. Statistical analysis: Mann–Whitney test. p < 0.05.

